# Administration of amniotic fluid stem cell extracellular vesicles promotes development of fetal hypoplastic lungs by immunomodulating lung macrophages

**DOI:** 10.1101/2022.11.29.518388

**Authors:** Lina Antounians, Rebeca Lopes Figueira, Bharti Kukreja, Elke Zani-Ruttenstock, Kasra Khalaj, Louise Montalva, Fabian Doktor, Mikal Obed, Matisse Blundell, Taiyi Wu, Cadia Chan, Richard Wagner, Martin Lacher, Michael D. Wilson, Brian T. Kalish, Augusto Zani

**Author notes:** These authors contributed equally to this work (Lina Antounians, Rebeca Lopes Figueira).

## Abstract

Congenital diaphragmatic hernia (CDH) is a devastating condition characterized by incomplete closure of the diaphragm and herniation of abdominal organs into the chest. As a result, fetuses have pulmonary hypoplasia, whose severity is the main determinant of poor outcome. The pathogenesis of pulmonary hypoplasia secondary to CDH is at least in part explained by lack or dysregulation of miRNAs that are known to regulate lung developmental processes. Herein, we report that intra-amniotic administration of extracellular vesicles derived from amniotic fluid stem cells (AFSC-EVs) rescues lung growth and maturation in a fetal rat model of CDH. To understand which fetal lung cells and biological pathways are affected by AFSC-EVs, we conducted whole lung single nucleus RNA-sequencing. We discovered that CDH lungs have a multilineage inflammatory signature with macrophage enrichment, and confirmed these findings in autopsy samples of lungs from human fetuses with CDH. Transcriptomic analysis of CDH fetal rat lungs also showed that AFSC-EV treatment reduced macrophage density and inflammation to normal levels. Analyzing the miRNAs contained in the AFSC-EV cargo with validated mRNA targets, we found that the downregulated genes in AFSC-EV treated CDH lungs were involved in inflammatory response and immune system processes. This study reports a single cell atlas of normal and hypoplastic CDH fetal rat lungs and provides evidence that AFSC-EVs restore lung development by addressing multiple pathophysiological aspects of CDH.

**One Sentence Summary:** Amniotic fluid stem cell extracellular vesicle treatment for fetal lung macrophage modulation

## Introduction

Pulmonary hypoplasia is a devastating condition characterized by impaired development of the fetal lung (*1*). A common cause of pulmonary hypoplasia is congenital diaphragmatic hernia (CDH), a defect due to incomplete closure of the diaphragm and herniation of abdominal organs into the chest (*2*). Hypoplastic lungs have fewer branches and alveoli, undifferentiated epithelium and mesenchyme, and vascular remodeling, characterized by fewer pulmonary vessels with muscularized wall layers and dysfunctional endothelium (*2, 3*). In babies with CDH, the severity of pulmonary hypoplasia is the main determinant of morbidity and mortality. In high-income countries, mortality has plateaued at 30% in the last three decades and can be as high as 90% in low- and middle-income countries (*4, 5*). Due to severe pulmonary hypoplasia, some fetuses die in utero or are electively terminated (*6*), some succumb in the first days of life without undergoing diaphragmatic repair (*4*), and many who survive and undergo surgery do not regain normal lung development (*2*). Longitudinal studies have shown that CDH survivors have long-term lung morbidity that persists beyond school age (*7, 8*). As there is consensus that the antenatal period offers a window of opportunity to reverse pulmonary hypoplasia, attempts have been made to promote fetal lung development antenatally (*9, 10*).

It is well known that lung developmental processes are regulated by multiple miRNAs (*11*), whose expression is missing or dysregulated in experimental and human CDH lungs (*12–24*). A promising avenue to deliver a heterogenous complement of miRNAs is based on the administration of extracellular vesicles (EVs). EVs are lipid-bound nanoparticles secreted by all cells for intercellular communication during physiological and pathological processes (*25*). EVs carry cargo in the form of genetic material (including miRNAs), proteins, and lipids, and transfer their cargo to target cells to induce biological responses (*25*). We previously reported that administration of EVs derived from amniotic fluid stem cells (AFSC-EVs) promotes branching morphogenesis, rescues tissue homeostasis, and stimulates epithelial cell and fibroblast differentiation in fetal rodent models of pulmonary hypoplasia (*12*). The ability to stimulate lung cell differentiation and rescue dysregulated signaling pathways was observed when AFSC-EVs were administered at canalicular and saccular stages of lung development, timepoints that are amenable to human translation (*26*). We performed enzymatic and inhibitor studies and found that the regenerative effects observed in hypoplastic lungs following AFSC-EV treatment were exerted at least in part via the release of their RNA cargo (*12, 27*). AFSC-EV RNA sequencing revealed that the cargo contained miRNAs that regulate the expression of genes involved in lung development, such as the miRNA 17~92 cluster (*12, 28*). However, it remains undetermined which lung cells are affected by AFSC-EVs and how AFSC-EVs rescue the biological pathways required for lung development. Herein, we used single nucleus RNA sequencing (snRNA-seq) to uncover the dysregulated genes and biological pathways in CDH fetal lungs and to determine the effects of in utero AFSC-EV therapy on fetal lung cell populations.

## Results

### Intra-amniotic administration of AFSC-EVs rescues branching morphogenesis and epithelial cell differentiation in fetal rats with CDH

To induce CDH, we used the experimental model based on the administration of nitrofen (2,4-Dichloro-1-(4-nitrophenoxy)benzene) to the rat dam at embryonic (E) day 9.5 (*29–32*). This is considered robust as it reproduces pulmonary hypoplasia in the whole litter with an analogous phenotype to the human condition (*29–32*). As the ideal time for fetal intervention in human babies with CDH has been identified as early as the canalicular stage of lung development (*33, 34*), we selected this stage in rats (*31*) to trial different routes of AFSC-EV administration. First, we opted for topical administration to fetal lungs via intra-tracheal instillation of AFSC-EVs. However, given the small size of fetal rats and the technical challenges with this survival surgery, we experienced a low rate of fetal survival (20%, n=10), as also reported by other groups (*35, 36*), and abandoned this route. We then tested two routes of administration that proved to have high fetal survival rates: intra-amniotic (IA, n=48, survival 84%) and maternal intra-venous (IV, n=30, survival 100%). When we compared the efficiency of AFSC-EV delivery to the fetus, we found that both IA and IV injection routes successfully delivered AFSC-EVs (fluorescently labeled with ExoGlow-Vivo) to fetal organs (Fig. 1A, fig. S1, and movie S1-6). However, we detected a positive fluorescent signal in fetal lungs only upon IA injection (Fig. 1B). To validate that the IA delivery of AFSC-EVs promoted lung growth and maturation in vivo, we assessed lung branching morphogenesis and cell differentiation markers at E21.5. Compared to control, CDH lungs had a reduction in airspace density and lower gene expression levels of lung maturation markers, such as fibroblast growth factor-10 (*Fgf10;* regulator of lung lineage commitment and branching morphogenesis), podoplanin (*Pdpn;* alveolar type 1 cell marker), and surfactant protein C and A (*Sftpc, Sftpa;* alveolar type 2 cell marker) (Fig. 1, C to E). CDH lungs from fetuses that received an IA injection of AFSC-EVs had restored airspace density and gene expression of *Fgf10, Pdpn, Sftpc*, and *Sftpa* back to control levels (Fig. 1, C to E). We validated these findings with immunofluorescence and Western blotting and determined that CDH lungs had reduced levels of PDPN and SPC compared to control (Fig. 1, F and G). Conversely, CDH lungs treated with AFSC-EVs had increased protein expression levels of PDPN and SPC.

**Fig. 1.**
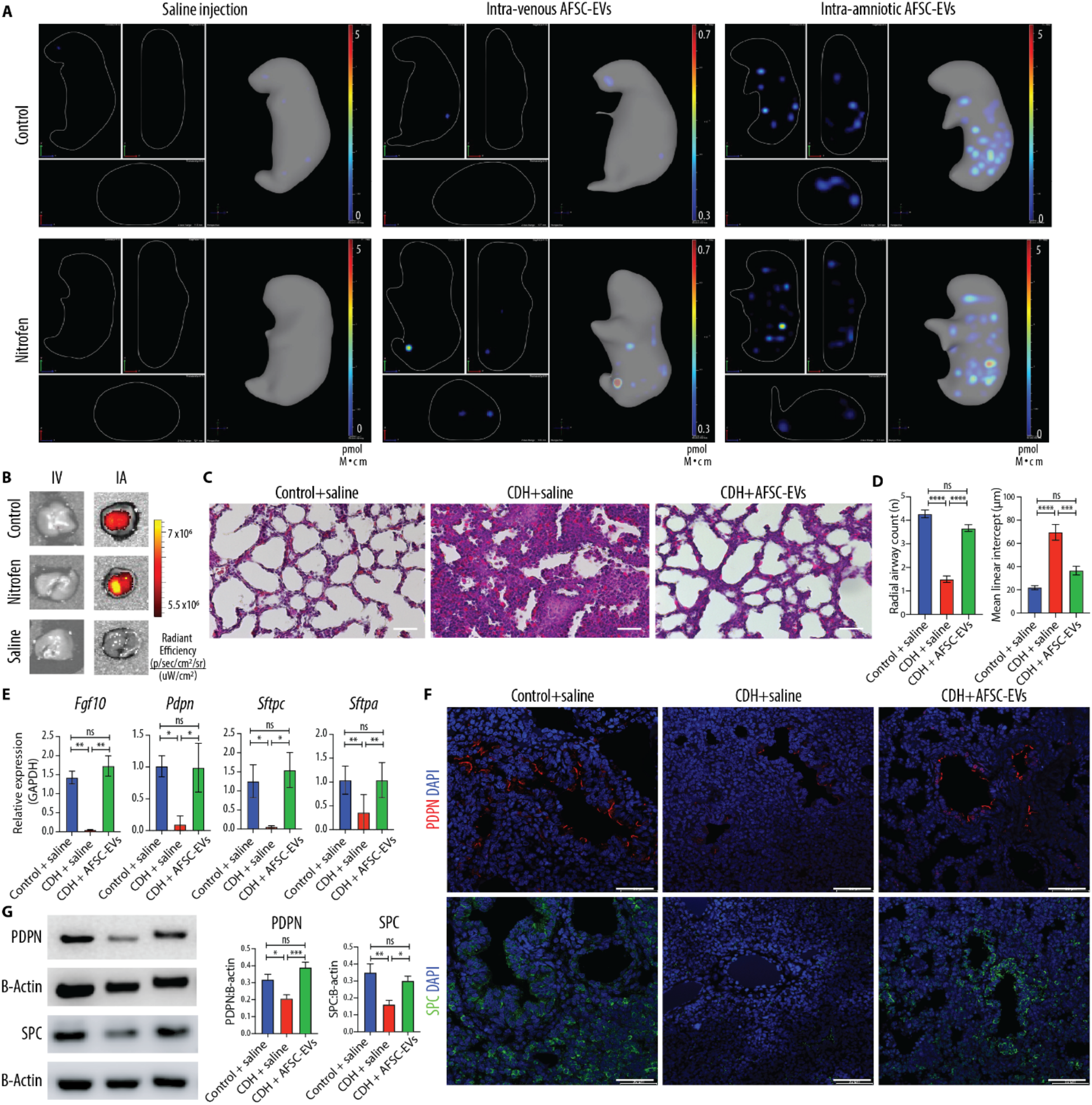
In vivo administration of AFSC-EVs reaches fetal lungs and rescues lung development in fetal rats with CDH. (**A**) Representative IVIS Spectrum instrument cross-sectional images from 3D bioluminescence reconstructions of whole fetuses at E21.5. Fetuses received either saline injection (left), intra-venous (IV) injection of ExoGlowVivo-stained AFSC-EVs (middle), or intra-amniotic (IA) injection of ExoGlowVivo-stained AFSC-EVs (right) at E18.5, in control fetuses (top row) or fetuses with pulmonary hypoplasia/CDH that received nitrofen (bottom row). Scale bar shows background-corrected fluorescence in pmol M^-1^ cm^-1^. Control+saline (n=3), Control+IV-AFSC-EVs (n=3), Control+IA-AFSC-EVs (n=7), Nitrofen+saline (n=6), Nitrofen+IV-AFSC-EVs (n=3), Nitrofen+IA-AFSC-EVs (n=16). No fluorescent signal was detected in saline injected controls. (**B**) Representative 2D optical images of dissected fetal lungs from the same conditions described in (A), quantified as radiant efficiency [p/s/sr]/[μW/cm^2^]. Control+saline (n=3), Control+IV-AFSC-EVs (n=3), Control+IA-AFSC-EVs (n=3), Nitrofen+saline (n=3), Nitrofen+IV-AFSC-EVs (n=4), Nitrofen+IA-AFSC-EVs (n=9). (**C**) Representative histology images (hematoxylin/eosin) of fetal lungs from Control+saline, CDH+saline, and CDH+AFSC-EV fetuses. Each condition included fetal lungs from n=5 experiments. Scale bar = 50 μm. (**D**) Differences in number of alveoli (radial alveolar count, RAC) in Control+saline (n=8), CDH+saline (n=8), CDH+AFSC-EVs (n=9), and paranchyme/airspace ratio (mean linear intercept, MLI) in Control+saline (n=8), CDH+saline (n=6), CDH+AFSC-EVs (n=6), quantified in at least 5 fields per fetal lung. ****P<0.0001, ***P<0.001. (**E**) Gene expression changes in lung developmental markers fibroblast growth factor-10 (*Fgf10*),podoplanin (*Pdpn*), a marker of alveolar type 1 (AT1) cells), and surfactant protein C (*Sftpc*) and A (*Sftpa*), a marker of alveolar type 2 (AT2) cells. Control+saline (n=5), CDH+saline (n=5), CDH+AFSC-EVs (n=5). **P<0.01, *P<0.05. (**F**) Representative immunofluorescence images of PDPN (red, top) and SPC (green, bottom) protein expression differences between Control+saline, CDH+saline, and CDH+AFSC-EVs fetuses (DAPI, blue). Scale bar = 50 μm. (**G**) Western blot analysis of PDPN and SPC expression in fetal lung quantified by signal intensity normalized to GAPDH. Control+saline (n=6), CDH+saline (n=7), CDH+AFSC-EVs (n=6). Groups were compared using Kruskal-Wallis (post hoc Dunn’s nonparametric comparison) for (D, RAC) and (E, *Pdpn* and *Sftpc*), Brown-Forsythe and Welch ANOVA (Dunnett’s T3 multiple comparisons test) for (E, *Fgf10*), and one-way ANOVA (Tukey post-test) for (D, MLI), (E, *Sftpa*), and (G), according to Shapiro-Wilk normality test.

### Single nucleus interrogation of the rat fetal normal and hypoplastic lung identifies four major cell types each with distinct subpopulations

To identify AFSC-EV cell-type specific effects, we conducted snRNA-seq on the left lung harvested at E21.5 from two IA saline injected controls, three IA saline injected left-sided CDH fetuses, and three IA AFSC-EV injected left-sided CDH fetuses (Fig. 2A). We selected fetuses for snRNA-seq studies from a large cohort of pups based on severity of branching morphogenesis and expression of lung maturation markers (Fig. 2B). After quality control filtering, we profiled a total of 298,653 nuclei (fig. S2 and Table S1). Analysis using Seurat revealed 15 distinct clusters representative of the four major cell types in the lung that corresponded to epithelial, endothelial, mesenchymal, and immune cells, each containing distinct subpopulations (Fig. 2, C and D). To the best of our knowledge, this is the first single-cell transcriptomic analysis of rat fetal lungs. Therefore, we used LungMAP, LungCellMap, and Human Protein Atlas annotations from mouse and human lungs to assign cell type identity based on gene expression enrichment of key marker genes (*37–39*). We identified five distinct epithelial sub-populations, including alveolar type I (AT1), alveolar type II (AT2), and ciliated epithelial cells (Fig. 2, C and D, and fig. S3). Among these cell types, AT1 cells expressed *Hopx, Pdpn, Clic5*, and *Ager;* AT2 cells expressed *Napsa, Lamp3, Fgfr2*, and *Etv5;* ciliated epithelial cells expressed cilia-related genes *Dnah12, Hydin, Ak9*, and *Spag17*. In addition, there were two other epithelial cell clusters with an inflammatory signature: cluster 8 was called “inflamed AT2 cells” as it co-expressed AT2 cell (*Lamp3, Lgi3*) and inflammatory markers, whereas cluster 14, broadly called “inflamed epithelial cells” expressed *Lcn2, Cc14, Cc16*, and *Il1b*. We identified two distinct endothelial clusters, one that had canonical endothelial cell markers *Tie2, Flt1*, and *Kdr*, and one that co-expressed mesenchymal and endothelial signatures (*Nfib, Tbx5, Adamts17*, and *Robo1*) that was termed “EndMT cells” (Fig. 2, C and D, and fig. S3). Five mesenchymal cell subtypes were identified, including fibroblasts, myofibroblasts, mesothelial cells, and pericytes (Fig. 2, C and D, and fig. S3). Among these cell types, fibroblasts expressed *Slit2, Fgf10*, and *Macf1*, myofibroblasts expressed *Myh11, Enpp2*, and *Pdgfra*, mesothelial cells expressed *Gpm6a, Aldh1a2*, and *Wt1*, and pericytes expressed *Pdgfrb, Ebf1*, and *Gucy1a1*. Moreover, we identified a mesenchymal cluster that heavily expressed *Lcn2, Ccl4, Cxcl1, Ccl3*, and *Plac8*, and was called “inflamed fibroblasts”. Lastly, three immune cell clusters were detected: two clusters expressing macrophage markers *CD68, Adgre1, CD163*, and *CD86*, and one expressing immune cell markers *Aoah, Zeb2*, and *Lyn* (Fig. 2, C and D).

**Fig. 2.**
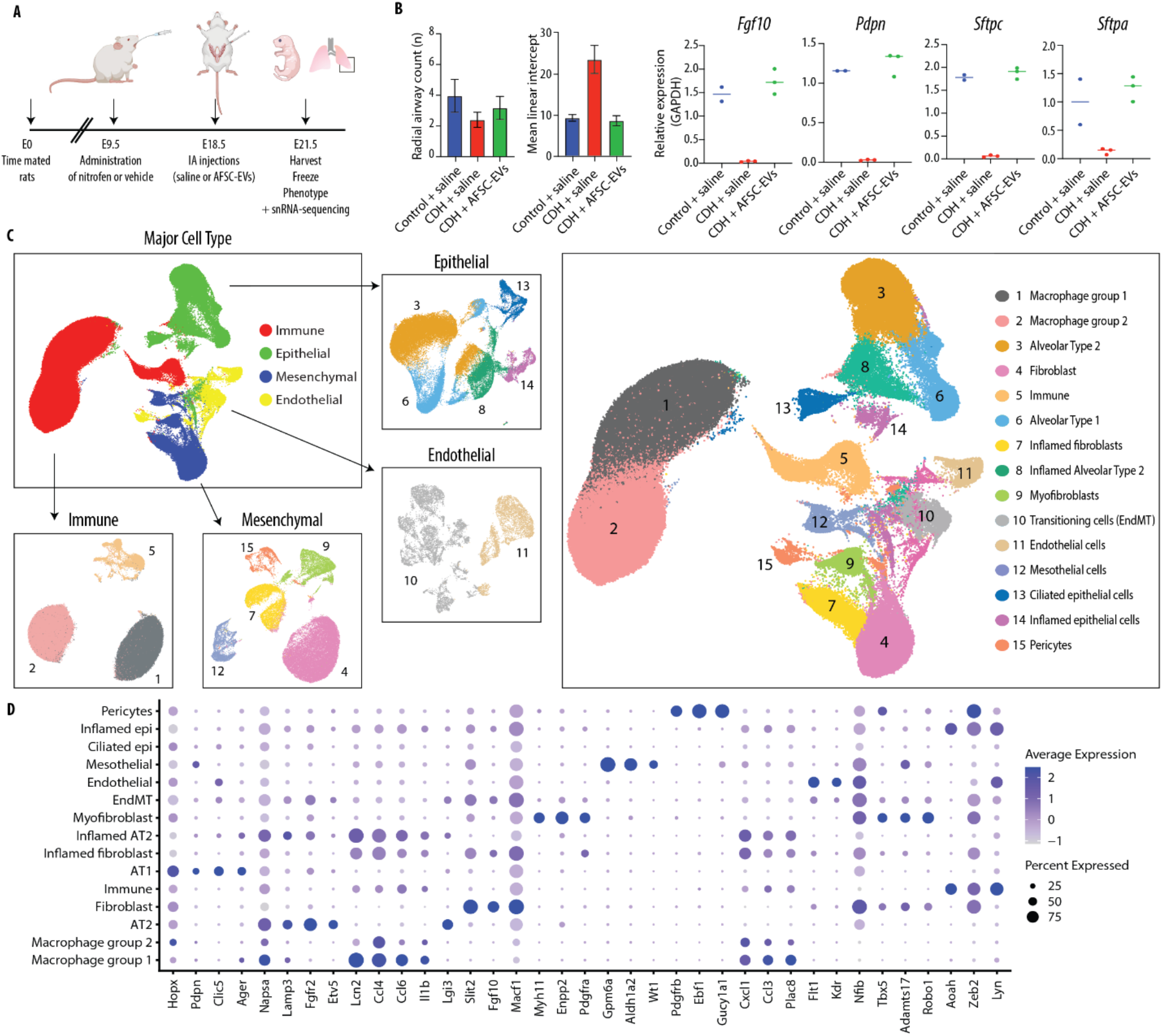
Single nucleus interrogation of the rat fetal normal and hypoplastic lung identifies four major cell types each with distinct subpopulations. (**A**) Schematic of experimental design and in vivo administration of AFSC-EVs in the rat model of CDH. (**B**) Criteria used for selection of 8 fetuses to undergo snRNA-seq analysis. Quantification of radial airspace count (RAC), paranchyme/airspace ratio (MLI), and gene expression differences in lung developmental markers from the three experimental groups. Control+saline (n=2), CDH+saline (n=3), and CDH+AFSC-EV (n=3). (**C**) Global UMAP projections of all nuclei (n=298,653) included in our study, further delineated by major cell type and subtype. (**D**) Expression of known cell type specific markers used to distinguish cellular subtypes within major cell type clusters. Node size is proportional to the percentage of nuclei within the specified cluster and node color denotes the average expression across nuclei within the specified cluster.

### Ligand-receptor analysis of rat fetal lung transcriptomics reveals the biological pathways that are influenced by AFSC-EV administration

To identify signaling pathways activated in CDH lungs treated with saline or AFSC-EVs, we performed ligand-receptor analysis on our snRNA-seq data using CellChat (*40*). We found that of all cell types, lung fibroblasts had the strongest outgoing signals and endothelial cells were the most receptive to incoming ligands (Fig. 3A). Our data also indicated that compared to normal lungs and AFSC-EV treated CDH lungs, CDH+saline lungs had upregulated ligand-receptor signaling from fibroblasts to endothelial cells (Fig. 3B). Our ligand-receptor analysis revealed that CDH lungs treated with saline exhibited signaling networks that are involved in inflammation and immune response, such as *Visfatin* (Fig. 3, C and D). Visfatin is a pro-inflammatory cytokine that potentiates TNFα and IL-6 production in human peripheral blood mononuclear cells and has been proposed as a biomarker for acute lung injury (*41–44*). Moreover, the strongest outgoing ligand signal in saline-treated CDH lungs was pleiotrophin (*Ptn*), a signaling molecule involved in lung development (*45*), which in our experiments was released from inflamed fibroblast and signaled to its receptors *Sdc2* and *Ncl* on multiple lung cell types (Fig. 3, C to E). PTN signaling was still present in AFSC-EV treated CDH lungs, but its receptor pair changed and was predominantly *Ptprz1*, a molecule responsible for regulating hematopoietic progenitor cell homing and retention (Fig. 3E) (*46*). Moreover, only AFSC-EV treated CDH lungs had activated signaling networks that control epithelial branching morphogenesis [*Fgf10-Fgfr2*, (*47, 48*)], surfactant synthesis and alveolar development [*Nrg2-Erbb4*, (*49*)], distal lung branching and alveologenesis [*Igf2-Igf1r* (*50, 51*)], anti-apoptotic processes [*Ptn-Alk* (*52*)], and angiogenesis [*Vegfa-Kdr* (*53*); Fig. 3, C to E, and Table S2].

**Fig. 3.**
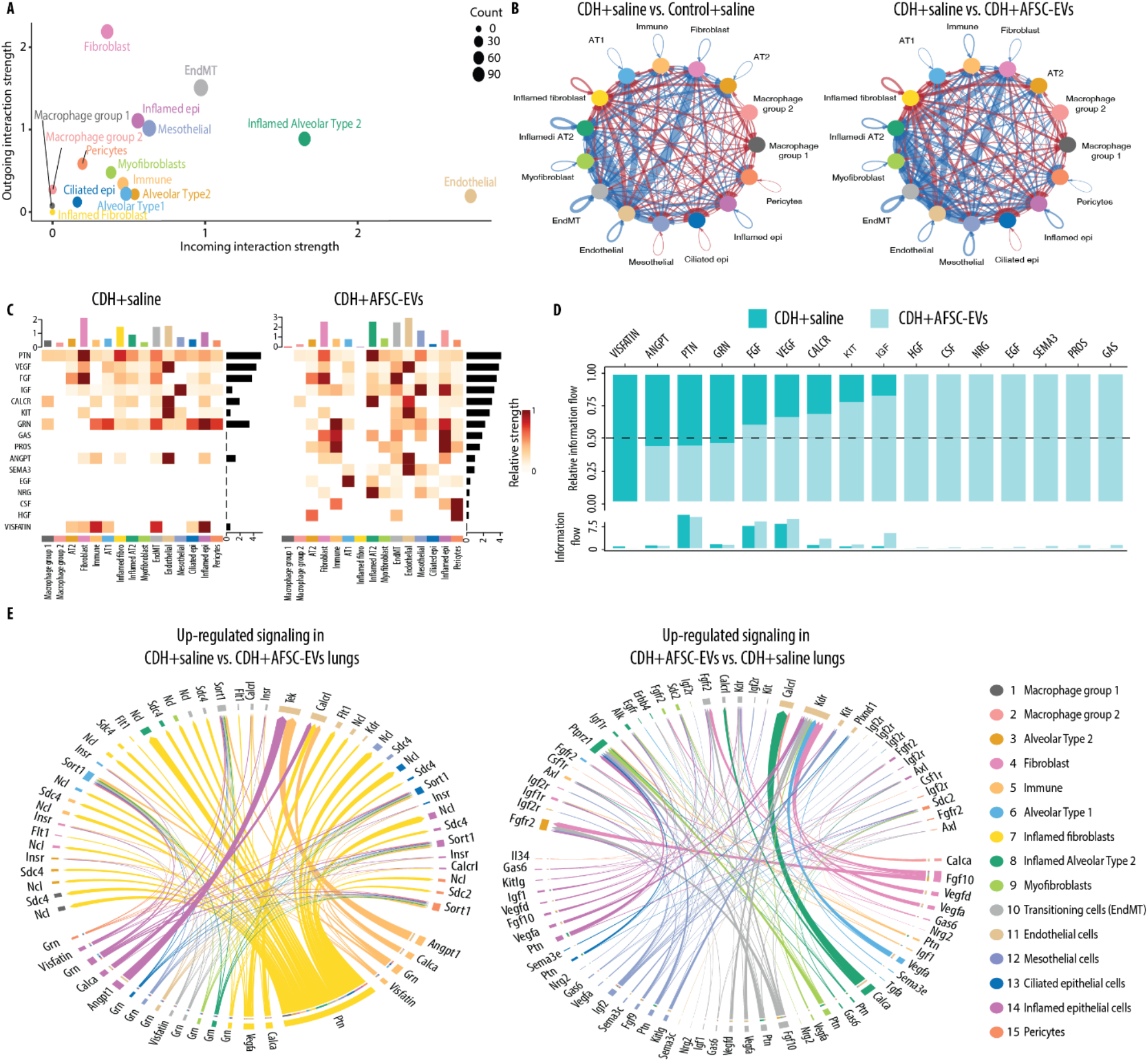
Ligand-receptor analysis reveals the biological pathways that are influenced by AFSC-EV treatment of rat fetal hypoplastic lungs. (**A-E**) CellChat analysis of signaling pathways in fetal lungs from all three conditions. (A) Comparison of interaction strength of outgoing and incoming signals by specific cluster. Node size represents number of interactions. (B) Significant interactions between clusters (arrows) showing number of interactions that are down-(blue) and up-regulated (red) when comparing Control+saline vs. CDH+saline (left) and CDH+saline vs. CDH+AFSC-EVs. Thickness of arrow indicates interaction strength. (C) Highly expressed ligand-receptor pairs displayed as a heatmap showing outgoing signal strength (top x axis), individual signaling pathways (left y-axis), strength of signaling pathway (right y-axis), and cell identity (bottom y axis). (D) Shift of signaling pathways related to lung development following AFSC-EV administration to fetal CDH lungs. (E) Chord diagram showing significantly up- or down-regulated signaling pathways in each cluster between CDH+saline and CDH+AFSC-EV conditions. Thickness of arrow indicates relative strength of specific pathway.

### CDH lungs have an inflammatory phenotype with high macrophage density that is rescued to normal levels by AFSC-EV administration

When we analyzed the snRNA-seq data by condition, we found that CDH+saline lungs had striking differences in the pattern and clustering of nuclei (Fig. 4A). Conversely, lungs from Control+saline and CDH+AFSC-EV groups had similar populations and distributions. We found that three clusters were unique to CDH+saline lungs, namely macrophage group 1 (cluster 1), inflamed fibroblasts (cluster 7), and inflamed ATII (cluster 8) (Fig. 4A). Moreover, macrophage group 2 was heavily represented in CDH+saline lungs (n=89,187 nuclei), compared to Control+saline (n=247 nuclei) and CDH+AFSC-EVs (n=739 nuclei; Table S1). Markers of macrophage identity and function were found in several clusters (Fig. 4B). To further delineate the specific macrophage subtypes contained in cluster 1 and 2, we used scPred, a validated machine-learning probability-based prediction method and trained the machine algorithm on single cell data from adult rat lungs (*54, 55*). We predicted that most nuclei in cluster 1 and 2 were from alveolar macrophages (n=140,382, 77%; fig. S4 and Table S3). Using immunofluorescence on the lungs of an additional cohort of rat fetuses, we confirmed a high density of macrophages in CDH+saline lungs, which was reduced to normal levels in CDH+AFSC-EV lungs (Fig. 4C).

**Fig. 4.**
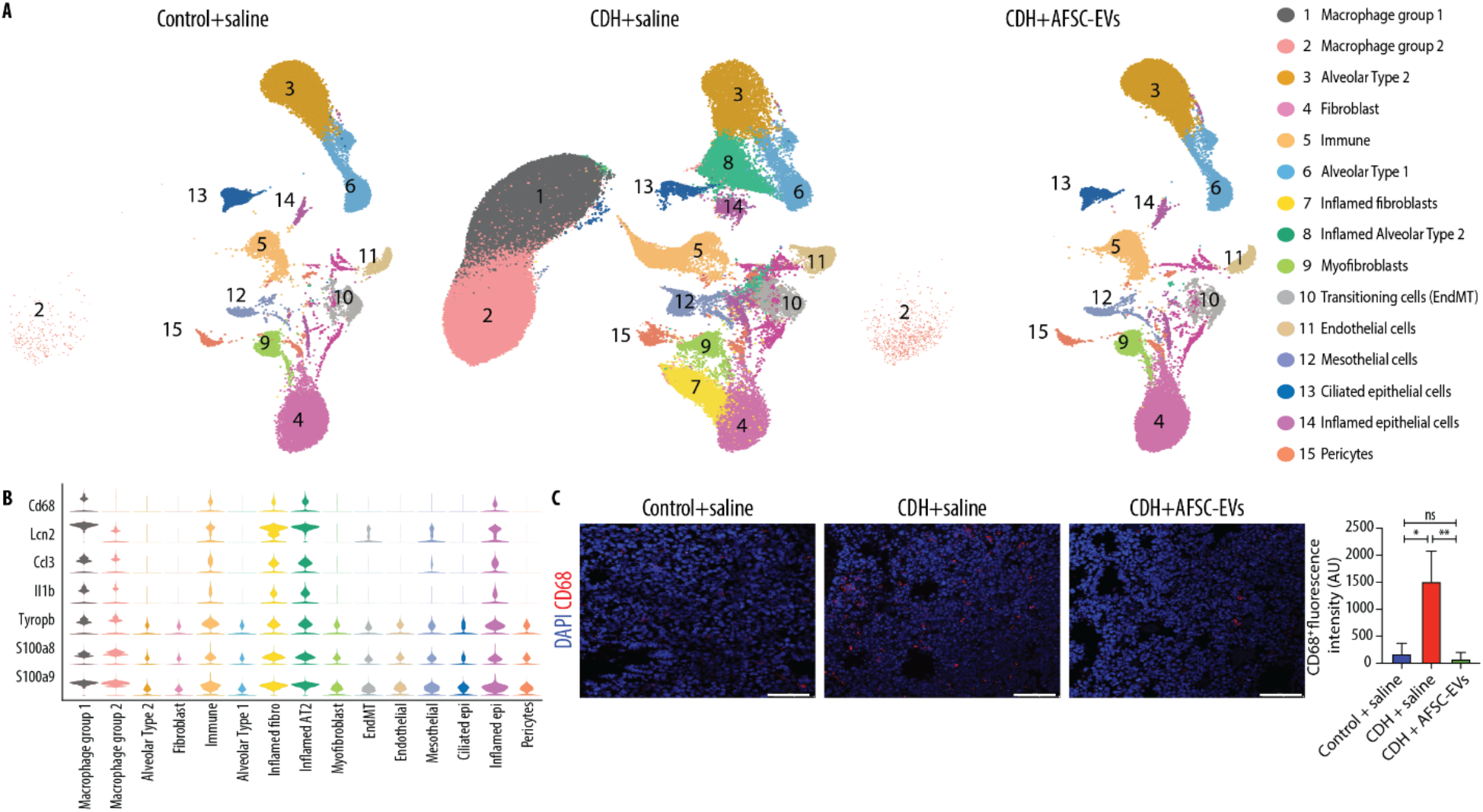
CDH lungs have an inflammatory phenotype with high macrophage density that is rescued to normal levels by AFSC-EV administration. (**A**) UMAP of snRNA-seq data split by condition. (**B**) Violin plots of macrophage and inflammatory marker gene expression across cell types, as measured by snRNA-seq. (**C**) Representative immunofluorescence images of panmacrophage marker CD68 in rat fetal lungs from all three conditions, quantified as fluorescence intensity of CD68 per field (AU, arbitrary units). Scale bar =50 μm. Control+saline (n=8), CDH+saline (n=6), and CDH+AFSC-EV (n=8). Groups were compared using Kruskal-Wallis (post hoc Dunn’s nonparametric comparison) for (C), according to Shapiro-Wilk normality test.

### CDH fetal lungs have a multilineage inflammatory signature that is dampened by the administration of AFSC-EVs

Differential gene expression analysis of CDH+saline lungs compared to Control+saline showed an extensive inflammatory signature across clusters with upregulation of *Il1b, Bcl2a1, Cxcl1, Ccl3/4*, and *Lcn2* (Fig. 5, A and B). These genes were downregulated in AFSC-EV treated lungs (Fig. 5, A and B, and Table S2). Differential gene expression analysis revealed similar patterns between Control+saline and CDH+AFSC-EV lungs regardless of the major cell type (Fig. 5C). Most of the highly differentially expressed genes in CDH+saline lungs were enriched for biological processes related to immune responses (Fig. 5D). Using immunofluorescence, we confirmed that the lungs of an additional cohort of CDH+saline rat fetuses were inflamed with upregulation of TNFα, which was restored to normal levels in those treated with AFSC-EVs (Fig. 5E).

**Fig. 5.**
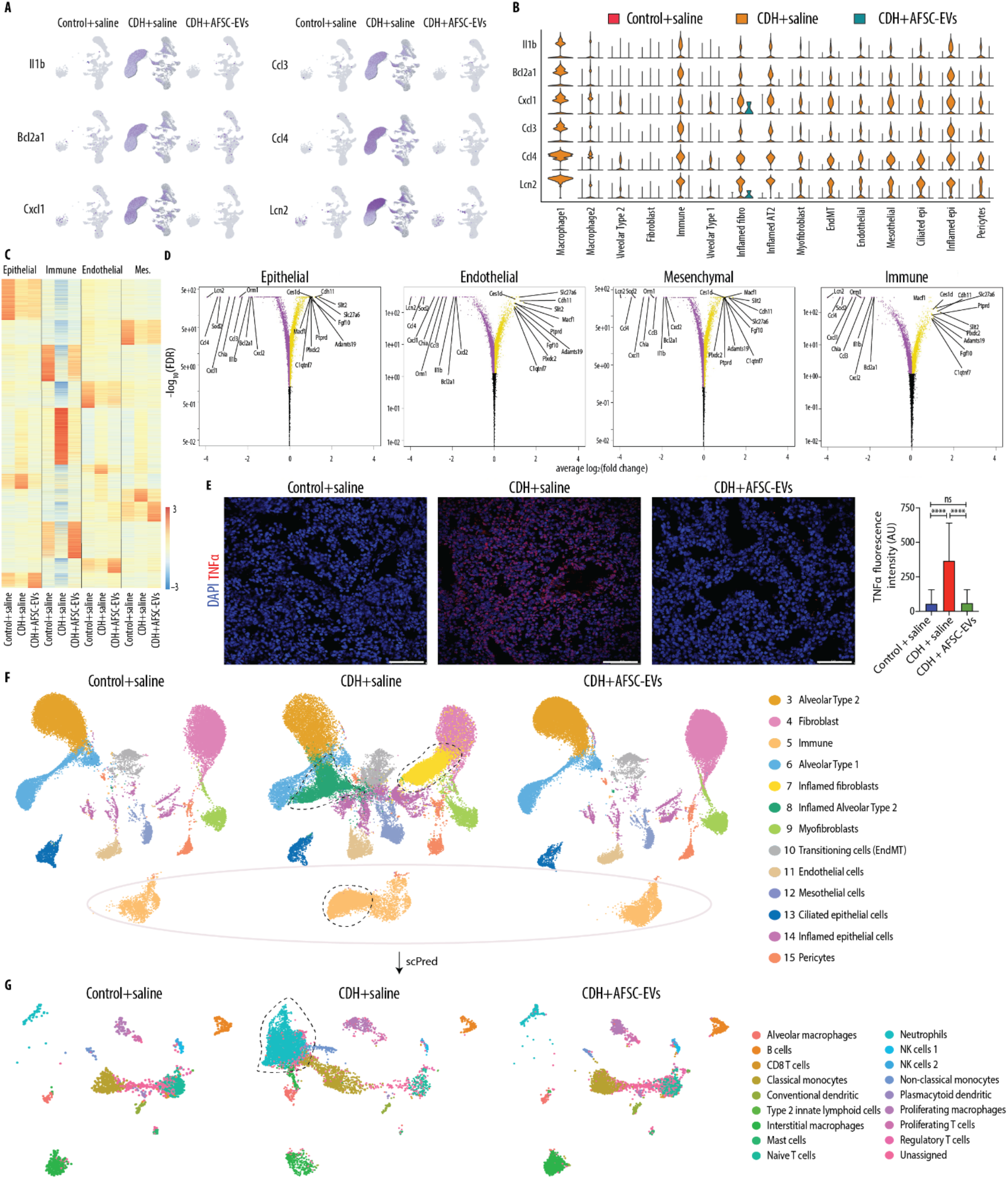
CDH fetal lungs have a multilineage inflammatory signature that is dampened by the administration of AFSC-EVs. (**A**) Featureplot of snRNA-seq data split by condition for six inflammatory genes with high expression in CDH+saline lungs. (**B**) Violin plot of inflammatory signature genes expression split by condition across cell types, as measured by snRNA-seq. (**C**) Heatmap displaying differential gene expression by major cell type, showing expression all genes ranked by log2-fold change and p-adjusted <0.05 within all conditions. (**D**) Volcano plots indicating most significantly differentially expressed genes by major cell type between CDH+saline and CDH+AFSC-EV treated groups. (**E**) Representative immunofluorescence images of inflammation marker TNFα in rat fetal lungs from all three conditions, quantified as density per mm^2^. Scale bar = 50 μm. Control+saline (n=5), CDH+saline (n=5), and CDH+AFSC-EV (n=5). (**F**) UMAP of a subset of data that excludes clusters 1 and 2 (overrepresented in CDH+saline group) split by condition. Outlines indicate nuclei or clusters that are represented in CDH+saline group compared to Control+saline and CDH+AFSC-EV groups. Control+saline (n=30,064), CDH+saline (n=45,114), CDH+AFSC-EV (n=42,193). (**G**) UMAP of predicted cell types contained in cluster 5 Immune cells from Fig. 5F, generated by machine learning algorithm (scPred) trained on rat adult lungs. Groups were compared using Kruskal-Wallis (post hoc Dunn’s nonparametric comparison) for (E), according to Shapiro-Wilk normality test.

To investigate the transcriptome differences across conditions and have a homogeneous comparison with similar number of nuclei within each condition, we created a subset of data by removing clusters 1 and 2 (macrophage group 1 and 2), as they were overrepresented in CDH+saline lungs. In this sub-analysis that included 30,064 Control+saline nuclei, 45,114 CDH+saline nuclei, and 42,193 CDH+AFSC-EV nuclei (fig. S2), we again found that all four major cell types were represented in all three conditions (Fig. 5F). Given the inflammatory signature of CDH+saline lungs, we investigated which immune cells were present in cluster 5 using scPred (*54, 55*). In all three conditions, we found different types of immune cells, including neutrophils, monocytes, and T and B cells (Fig. 5G, and Table S4). Among these immune cells, CDH+saline lungs had higher proportion of neutrophils compared to Control+saline lungs (53% vs. 3%, p<0.0001; Fisher’s exact test). Conversely, CDH+AFSC-EV lungs had a lower proportion of neutrophils (4%) compared to CDH+saline lungs (p<0.0001, Fisher’s exact test).

### Predicted miRNA-mRNA signaling pathways activated by AFSC-EVs

As we have shown that the regenerative effects of AFSC-EVs are mainly due to the delivery of miRNAs to fetal lung cells (*12, 27*), we used publicly available datasets (TarBase, MicroCosm, miRanda, miRDB, miRecords, miRTarBase) to generate a network between the miRNAs that are known to be present in the rat AFSC-EV cargo (*12*), and the mRNAs identified by snRNA-seq that were downregulated in CDH+AFSC-EV lungs compared to CDH+saline lungs. We found that several mRNAs that were downregulated in CDH+AFSC-EV lungs were regulated by multiple miRNAs that were present in the AFSC-EV cargo (Fig. 6, A and B). Overall, we found 820 predicted miRNA-mRNA targets that regulate several biological processes, including inflammatory/immune responses (Fig. 6C). From the 820 predicted miRNA-mRNA pairs, 32 miRNA-mRNA pairs (13 miRNAs, 24 mRNAs) have previously been validated (Fig. 6D).

**Fig. 6.**
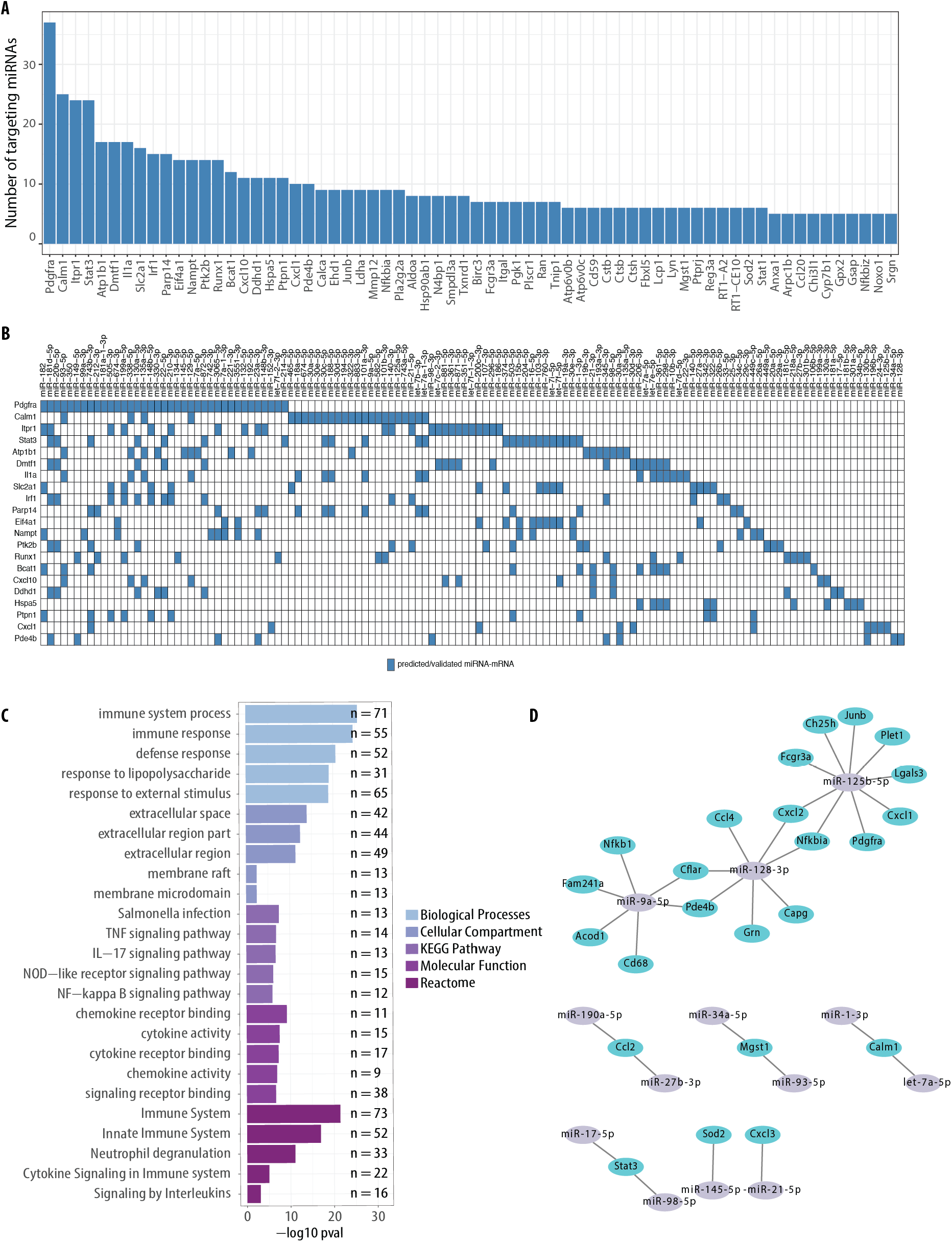
Predicted miRNA-mRNA signaling pathways activated by AFSC-EVs. (**A**) Bar graph indicating number of miRNA-mRNA interactions (y-axis) between AFSC-EV cargo miRNAs and down-regulated genes in CDH+AFSC-EV lungs (x-axis). Predicted miRNA-target interactions obtained from publicly available datasets (tarbase, MicroCosm, miRanda, miRDB, miRecords, miRTarBase). (**B**) Heatmap of specific AFSC-EV miRNAs (x-axis) and their redundant roles in downregulating genes in CDH+AFSC-EV lungs (y-axis). (**C**) Gene set enrichment analysis of the downregulated genes using g:Profiler. (**D**) Network of validated miRNA-mRNA pairs showing downregulated genes (blue nodes) and AFSC-EV miRNAs (purple nodes).

### Inflammatory markers are upregulated in hypoplastic lungs of human fetuses with CDH

To confirm that the findings observed in fetal rats are relevant to the human CDH condition, we interrogated lung sections from autopsy samples of four human fetuses with CDH that died between gestational weeks 19 and 26 (canalicular stage of lung development) and four controls (no fetal lung pathology or systemic inflammatory conditions, no chorioamnionitis; Table S5). We first confirmed that compared to controls, the lungs of the CDH fetuses had a lower density of airspaces (Fig. 7A). We then determined that the macrophage density was reduced in lungs of CDH fetuses, most predominantly in the parenchyma (Fig. 7B). Moreover, the expression of canonical markers of inflammation such as TNFα and its downstream NF-κB signaling (*56*) were upregulated in lungs of CDH fetuses (Fig. 7C).

**Fig. 7.**
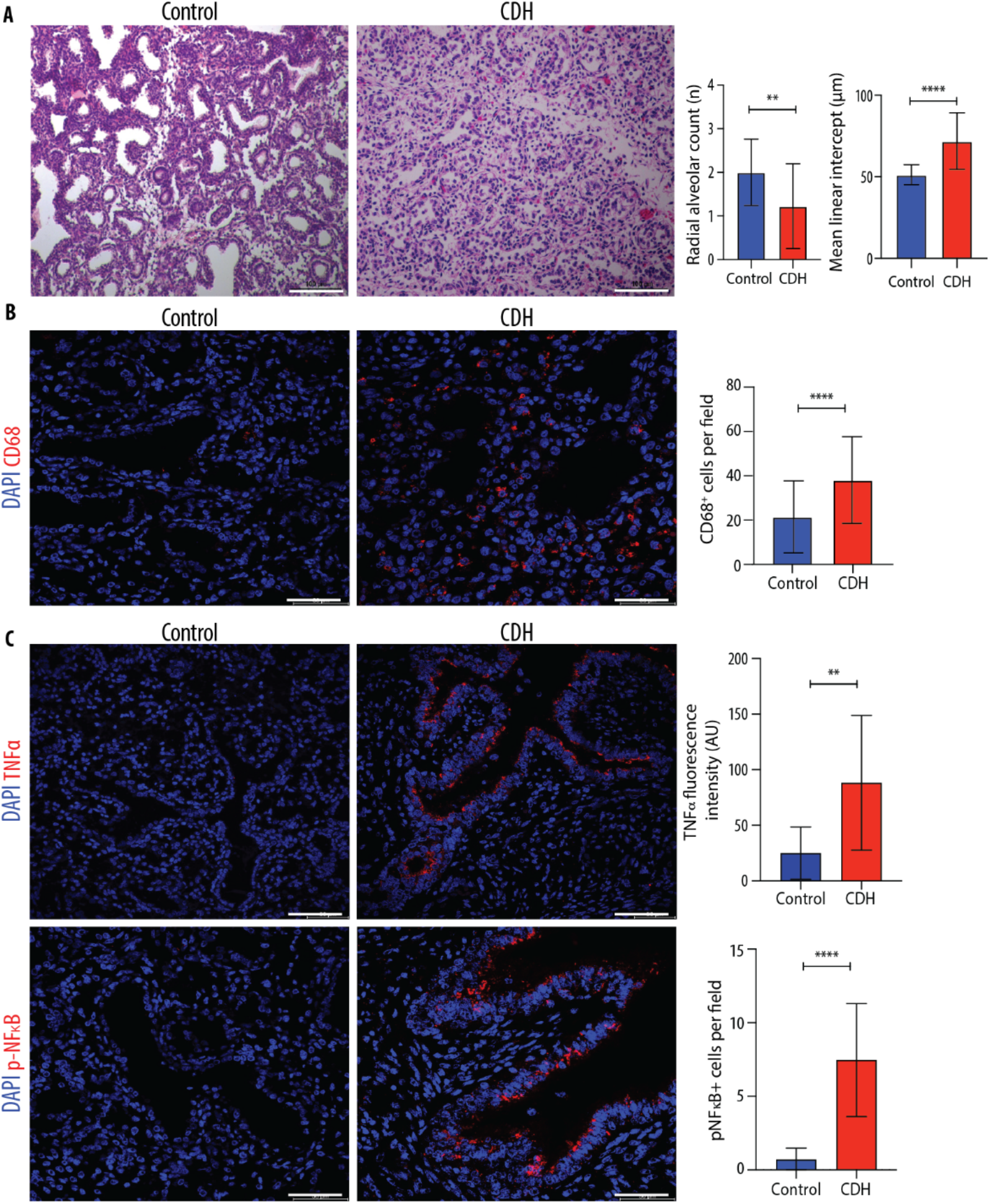
Hypoplastic lungs of human fetuses with CDH have increased macrophage density and upregulation of inflammatory mediators and can be targeted with human AFSC-EV treatment. (**A**) Representative histology images (hematoxylin/eosin) of fetal lungs from autopsy studies of CDH fetuses (n=4) and controls with no lung pathology or inflammatory condition (n=4). Scale bar = 100 μm. Quantification of paranchyme/airspace ratio (MLI) and number of alveoli (RAC) in 10 fields per fetal lung. (**B**) Representative immunofluorescence images of pan-macrophage marker CD68 in human fetal lungs autopsy samples from CDH (n=4) and controls (n=4), quantified as number per field. Scale bar = 50 μm. (**C**) Representative immunofluorescence images of inflammatory mediators TNFα and phosphorylated NF-κB (p-NF-κB) in the same experimental groups as (B) quantified by fluorescence intensity of TNFα and density of p-NF-κB^+^ cells per field. Scale bar = 50 μm. Groups were compared using two-tailed Mann Whitney test for (A), (B), and (C, pNF-κB), and two-tailed Student’s t-test for (C, TNFα), according to Shapiro-Wilk normality test.

## Discussion

Hypoplastic lungs of fetuses with CDH are classically described as having impaired growth (fewer branches and airspaces), maturation (undifferentiated epithelium and mesenchyme), and vascularization (fewer and muscularized lung vessels that undergo vascular remodeling) (*2*). We employed a transcriptomic approach at a single cell resolution in an experimental model of CDH and discovered that hypoplastic lungs also have an inflammatory signature with high density of macrophages and upregulation of biological pathways that are involved in inflammatory and innate immune response. Using immunofluorescence on autopsy samples, we confirmed that human fetuses with CDH also have an inflammatory status with macrophage enrichment and increased TNFα and pNF-κB expression. A similar observation was made in neonates with CDH postnatally, who were found to have upregulation of pNF-κB in the proximal lung (basal cells) and TNFα in the distal lung (*57, 58*). Other studies reported that neonates with CDH had high levels of proinflammatory cytokines in the blood postnatally (*59–64*). Moreover, using the nitrofen model, some studies have reported that rat fetuses with CDH had lung inflammatory changes such as high levels of monocyte chemoattractant protein-1 (*Mcp1;* also known as chemokine C-C motif ligand 2, *Ccl2*) and *Tnfα* (*65–67*). A recent study employing unbiased proteomics on the lungs of rat fetuses with CDH showed elevated levels of inflammatory mediators such as STAT3 compared to non-CDH controls (*68*). Our snRNA-seq experiments showed that all four major lung cell types in CDH fetuses had upregulation of several inflammatory mediators, including *Tnfa, Stat3*, and chemokines *Ccl3/4* and *Cxcl1*. Moreover, we report upregulation of pro-inflammatory mediators, such as lipocalin-2 (*Lcn2*), an acute-phase protein involved in the immune response to lung inflammation (*69*), and interleukin 1b (*Il1b*), a factor known to stimulate the immune response and cause disruption of lung morphogenesis (*70, 71*).

In our study, the multilineage inflammatory response was accompanied by an increase in macrophage density in the lung. Although there is robust literature on the role of macrophages in the adult lung during injury and repair (*72–75*), less is known about macrophage involvement in impaired perinatal lung development. In bronchopulmonary dysplasia (BPD), a condition typical of premature babies characterized by hypoplastic lungs with fewer branches and alveoli similar to CDH (*76, 77*), inhibition of branching morphogenesis is partly caused by the activation of fetal lung macrophages, and depletion or targeted inactivation of macrophages is protective against impaired branching morphogenesis (*78*). The role of macrophages in fetal hypoplastic CDH lungs remains undetermined. Since macrophages have different functions that range from pro-inflammatory to reparative, those recruited or activated in CDH fetal lungs could either contribute to the arrest in lung development as in BPD, or aid in the rescue of branching morphogenesis (*78–80*).

Given the multilineage inflammatory signature observed in CDH hypoplastic lungs, it is reasonable to think that inflammation plays a key role in the pathogenesis of pulmonary hypoplasia and contributes to poor prognosis in babies with CDH. Moreover, lung inflammation may provide an alternative target to rescue lung development. In the past, attempts have been made to promote lung maturation in CDH via prenatal administration of corticosteroids, whose anti-inflammatory effects are well known (*81, 82*). The rationale was based on promoting lung development by stimulating surfactant protein C production, as shown in several studies on premature lungs of neonates with BPD (*81, 83*). Although experimental studies in the nitrofen model showed an improvement of fetal lung maturation upon corticosteroid administration (*84–88*), large database retrospective studies and prospective clinical trials in CDH showed no benefit in survival, length of stay, ventilator days, or oxygen use at 30 days (*89, 90*). As a result, antenatal corticosteroid administration is not recommended as a standard practice in the care of patients with CDH (*89, 90*).

A promising avenue for targeting multiple molecules and pathways is through an EV-based therapy. We have previously shown that antenatal AFSC-EV administration rescues dysregulated signaling pathways relevant to lung development, and results in improvement of lung branching morphogenesis and epithelial and mesenchymal maturation (*12, 26*). In the current study, we provide evidence that AFSC-EVs also have anti-inflammatory effects in a robust fetal rat model of CDH. The anti-inflammatory effects of stem cell-derived EVs have been recognized within the last decade in numerous experimental and clinical trials in several conditions, including BPD (*91–93*). Furthermore, with the advent of SARS-CoV2-induced acute respiratory distress syndrome (ARDS), many research groups have attempted to use EVs from different sources, including stem cells, as a possible strategy to treat lung inflammation (*94, 95*). One clinical trial testing the efficacy of stem cell-derived EVs on reduction of ARDS symptoms reported that EV administration is safe, restores oxygenation, downregulates cytokine storm, and reconstitutes immunity in patients with ARDS (*96*). Evidence that stem cell-derived EVs have a beneficial effect also in premature lungs has been shown in several experimental studies (*97–104*). In these studies, mesenchymal stem/stromal cell-derived EVs (MSC-EVs) administered to different models of experimental BPD showed improvements in lung function, reversal of lung vascular remodeling and fibrosis, and attenuation of lung inflammation. MSC-EV beneficial effects to BPD lungs were found to be due to the epigenetic and phenotypic reprogramming of myeloid cells (*105*) and modulation of lung macrophage phenotype (*102*). While we had previously shown that MSC-EV administration partially promoted epithelial cell homeostasis and differentiation in CDH lungs, we did not observe a complete rescue of biological pathways and key phenotypic aspects of lung development, such as branching morphogenesis and alveolarization (*12, 26*). Conversely, we found that AFSC-EVs were a better candidate than MSC-EVs for reversing key features of pulmonary hypoplasia in CDH, likely due to their RNA cargo that was enriched with miRNAs responsible for lung developmental processes at a comparative analysis (*12*). In the present study, we have shown that AFSC-EVs rescue the density of macrophages and the expression of inflammatory mediators in the lung back to control levels. This observation is in line with a recent study that reported the ability of AFSC-EVs to modulate inflammasome activation in monocytic cells in vitro (*106*). Although immune cells are not predominant in fetal lungs, our snRNA-seq analysis was able to detect gene expression differences in immune cell populations. Moreover, several genes identified by our snRNA-seq analysis to be upregulated were also observed in a single cell RNA-sequencing study postnatal lungs of mice with BPD (*107*). This study found that upregulation of inflammatory cytokine signaling was associated with major structural and cell-to-cell signaling changes in the lung (*107*).

Although we provide evidence that AFSC-EV administration has potential for reversing pulmonary hypoplasia in rat fetuses with CDH, there are several steps that still need to be taken before translating this promising approach to clinical application, including the establishment of the optimal route of administration. In this study, we initially tested three routes and ultimately opted for AFSC-EV intra-amniotic injection during the saccular stage of lung development. At this timepoint, clustered fetal breathing movements occur at ~40 movements/hour in fetal rats (*108*) and intra-amniotically injected products reach the fetal lung, as reported (*27, 109–111*). Translating this approach to humans is feasible, as access to the amniotic sac is routine during pregnancy for diagnostic procedures, such as amniocentesis. Moreover, a study has reported that intra-amniotic administration of ectodysplasin A to human twins with X-linked hypohidrotic ectodermal dysplasia resulted in a profound reversal of their disease phenotype (*112*). Herein, we also tested the intra-tracheal route, which had a high rate of fetal demise due to the invasiveness of the procedure in fetal rodents, as reported (*35, 36*). Nonetheless, this route could be tested in a larger animal model, such as the lamb, and possibly considered to be used in conjunction with fetal endoscopic tracheal occlusion (FETO) in human babies. FETO is a surgery procedure based on the deployment of a balloon in the fetal trachea to prevent egression of amniotic fluid and promote lung growth (*2*). Two randomized controlled trials reported 25% improved survival following FETO in severe but not moderate CDH (*113, 114*). Lastly, we tested the maternal intra-venous route of administration and demonstrated that AFSC-EVs cross the placental barrier, a property of EVs that has been well-described (*115, 116*). Although this strategy would circumvent the invasiveness of fetal intervention for both mother and fetus, efforts should be focused on custom designed EVs that could specifically target the fetal lung and avoid off-target effects. Alternatively, the fetal circulation can be directly accessed through the umbilical vein as recently shown in a fetus with Pompe’s disease (*117*). In this case, the fetus received multiple infusions of in utero enzyme-replacement therapy, administered under ultrasonic guidance from 24 to 34 weeks of gestation, and resulted in reversal of cardiac and motor function deficits (*117*).

We acknowledge that our study has some limitations. Due to the poor capture of lowly expressed transcripts, our snRNA-seq analysis could not be used to discern the origin (monocyte-derived vs. tissue resident) or the polarization status (M1-like pro-inflammatory vs. M2-like reparative) of CD68^+^ cells. Moreover, as snRNA-seq transcriptomics requires tissue dissociation, we were not able to define where the CD68^+^ cells were located within the lung, thus not being able to discern between alveolar and interstitial macrophages. Nonetheless, using one other adult rat lung single cell RNA dataset, we were able to make predictions on which types of macrophages reside in our sequencing clusters. Furthermore, without conducting genetic knock-down studies, it is difficult to ascertain if macrophages are at the root cause of the pathogenesis of pulmonary hypoplasia secondary to CDH or if they are recruited and activated by an increased inflammatory state. Similarly, it remains unclear how AFSC-EVs induce a reduction in macrophage density. Further studies are underway to address these important questions before translating these findings to human patients with CDH.

## Materials and Methods

### Study design

The objective of this study was to investigate the AFSC-EV regenerative ability and mechanism of action on fetal hypoplastic lungs through in vivo administration and snRNA-seq. The rat model of CDH and pulmonary hypoplasia was used for part of this study, as obtaining fresh human CDH lung tissue is not considered ethically acceptable. Fetal rat lungs with CDH closely resemble the degree of pulmonary hypoplasia that is observed in human fetuses with CDH. Findings were confirmed on lung autopsy samples obtained from four fetuses with CDH and appropriate controls. As detailed below, experimental models and sample collection were approved by the regulatory committee at The Hospital for Sick Children, Toronto (AUP#49892 and REB#1000074888). Sprague-Dawley rats were randomly assigned to treatment groups. All data including outliers is shown, and all experiments were performed in at least triplicate, with the number of replicates indicated in the figure legends. Additional details on the methods used in this study are provided in the Supplementary Materials.

### EV isolation, characterization, and tracking

EVs from rat AFSCs were isolated and characterized as described previously (*12*). Briefly, rat AFSC conditioned medium was obtained by treating cells with exosome-depleted FBS for 18 hours. EVs were isolated by differential ultracentrifugation as previously described (*118*). Rat AFSC-EVs were previously characterized in accordance with the International Society for Extracellular Vesicles guidelines for proper size, morphology, and expression of canonical EV protein markers (*12*). To track AFSC-EV migration in vivo, ExoGlowVivo™ was used following the manufacturer’s recommended protocol, with 250 μg protein equivalent of AFSC-EVs stained using 2 μL of ExoGlow™-Vivo (Near IR) EV Labeling Kit (System Biosciences, Palo Alto, CA).

### Experimental model of pulmonary hypoplasia

Pulmonary hypoplasia was induced in rat fetuses as previously described (*29–32*) with the administration of 100 mg nitrofen to pregnant rat dams by oral gavage on E9.5. For in vivo AFSC-EV administration, three routes were used and described in detail in the Supplementary Materials. The IA route of administration was conducted by anesthetizing rat dams at E18.5, exposing uterine horns through a midline laparotomy, and injecting 100 μL of AFSC-EVs intra-amniotically with a 30G needle, away from the body but close to the face of the fetus. Rat dams and fetuses were monitored in accordance with Canadian Council on Animal Care guidelines. At E21.5, rat dams were anesthetized, uterine horns exposed, and each fetus was examined for survival by assessing fetal size and movement. Fetuses were then delivered, immediately euthanized, and fetal lungs were perfused with saline to remove red blood cells then immediately frozen or placed into 4% paraformaldehyde solution. Samples were stored at −80 °C for RNA/protein analysis or at 4 °C for histology.

### Outcome measures

For EV tracking, ExoGlow™-Vivo labelled AFSC-EVs (784 nm excitation, 820 nm emission) were injected through IA or IV route of administration. Whole fetuses, individual fetal organs, and maternal organs were imaged using the IVIS^®^ Spectrum In Vivo Imaging System – PerkinElmer (CFI Facility, University of Toronto). For lung morphometry, lung sections of fetal rats or human autopsy samples were stained with hematoxylin and eosin, and analyzed for radial airspace count and mean linear intercept, as previously described (*12, 27*), and recommended by the American Thoracic Society (*119*). For assessment of lung development, gene expression analysis was conducted using quantitative polymerase chain reaction (RT-qPCR), and protein expression analysis of SPC and PDPN was conducted using immunofluorescence assays and Western blotting.

To study the transcriptomic changes at the single cell level, a subset of fetal lung samples was chosen (Control+saline, n=2; CDH+saline, n=3; and CDH+AFSC-EVs, n=3), and nuclei were extracted and subjected to the 10X Genomics protocol (NovaSeq 6000). Data files were obtained with Cellranger-6.0.0, aligned to the rattus norvegicus version 6 genome, and analyzed using Seurat (4.0.3). Differential gene expression was determined with FindMarkers (MAST), and ligand-receptor analysis was performed using CellChat (1.1.3, *40*) with default parameters. Further details on sequencing analysis are described in supplementary materials. To assess the presence of macrophages, immunofluorescence staining of pan-macrophage marker CD68 was used on fetal rat lungs and human lung autopsy samples. Immunofluorescence assays of inflammatory mediators was also conducted to determine whether fetal rat lungs and human lung autopsy samples were inflamed. For human autopsy samples, two independent researchers differentiated between red blood cells and macrophages to assess CD68^+^ cells. To identify miRNA-mRNA regulatory pathways, AFSC-EV miRNA cargo analysis (*12*) and downregulated genes from snRNA-seq dataset were analyzed using multiMiR (version 1.16.0, *120*). Gene set enrichment analysis was conducted using g:Profiler (version 0.7.0, *121*).

### Statistical analysis

Experimental groups were compared using two-tailed Student’s t-test, Mann-Whitney, Fisher’s exact test, one-way ANOVA (Tukey post-test), Kruskal-Wallis (post-hoc Dunn’s nonparametric comparison), or Brown-Forsythe and Welch ANOVA (Dunnett’s T3 multiple comparisons test) tests according to Gaussian distribution assessed by Shapiro-Wilk normality test. P value <0.05 was considered significant. All statistical analyses were produced using GraphPad Prism^®^ software version 6.0. Differential gene expression analysis was conducted using BioConducter R (3.15) package MAST (1.22.0) between two conditions, with adjusted p-value <0.05 and log2(fold change) > |0.5| considered as significant for snRNA-seq experiments.

## List of Supplementary Materials

Movie S1. Representative 3-dimensional image of in vivo AFSC-EV tracking experiments for Control+AFSC-EVs.

Movie S2. Representative 3-dimensional image of in vivo AFSC-EV tracking experiments Nitrofen+AFSC-EVs.

Movie S3. Representative 3-dimensional image of in vivo AFSC-EV tracking experiments Nitrofen+saline (negative control).

Data file S1. Images and quantification of in vivo AFSC-EV administration in whole fetuses.

Data file S2. Images and quantification of in vivo AFSC-EV administration in fetal organs.

Data file S3. Data for Western blotting experiments.

Data file S4. Differentially expressed genes between Control+saline, CDH+saline, and CDH+AFSC-EVs by major cell type in fetal rat lungs.

Data file S5. Gene set enrichment analysis of differentially expressed genes between Control+saline vs. CDH+saline and CDH+saline vs. CDH+AFSC-EVs

MDAR Reproducibility Checklist

## Acknowledgements

The authors would like to thank G. Raffler, M. S. Gaffi, S. Gandhi, H. Hou, and A. Hardy. We are indebted to J. Reyes, D. Chiasson, and G. Somers for selection of autopsy samples included in the manuscript, P. De Coppi for providing rat AFSCs in kind, and the Lab Animal Service core facility at the Hospital for Sick Children. Some of the equipment used in this study was supported by the 3D (Diet, Digestive Tract and Disease) Centre funded by the Canadian Foundation for Innovation and Ontario Research Fund, project number 19442 and 30961.

## Author Contributions

Conceptualization: LA, RLF, BK, MDW, BK, AZ

Methodology: LA, RLF, EZR, KK, LM, CC, MDW, AZ

Visualization: LA, RLF, BK, EZR, KK, LM, FD, MO, MB, TW, CC, BK, AZ Funding acquisition: LA, RLF, KK, RW, ML, MDW, BK, AZ

Project administration: LA, AZ

Supervision: MDW, BK, AZ

Writing – original draft: LA, AZ

Writing – review & editing: LA, RLF, BK, CC, MDW, BK, AZ

## Data and materials availability

snRNA-sequencing data are available in the NCBI GEO (no. GSE211914). Relevant data for EV isolation and characterization was previously submitted to EV-TRACK knowledgebase (no. EV190001). RNA-sequencing for AFSC-EV cargo sequencing was accessed from ArrayExpress database (no. E-MTAB-8921) from our previous publication (*12*).

