## Supplementary Materials for "Administration of amniotic fluid stem cell extracellular vesicles promotes development of fetal hypoplastic lungs by immunomodulating lung macrophages"

**Extended Materials and Methods:**

*In vivo AFSC-EV administration routes*

Three routes of administration were used to administer AFSC-EVs in vivo in rats 1) intra-tracheal: dams were anesthetized at E18.5, uterine horns were exposed through a midline laparotomy, and the head of the fetus was exposed through a prolene 6.0 purse string suture. The fetal trachea was dissected and intra-tracheal injection of 5 µL of AFSC-EVs was conducted with a 30G needle. The fetus was then returned to the amniotic sac and the purse string suture was closed. 2) intra-amniotic: dams were anesthetized at E18.5, uterine horns were exposed through a midline laparotomy, and intra-amniotic injections were conducted with a 30G needle, away from the body but close to the face of the fetus. No sutures were required, and sacs were returned into the dam. 3) maternal intra-venous: dams were anesthetized at E18.5, tail vein was identified and AFSC-EVs were injected with a 25G needle. Samples were harvested at E21.5 for all three routes of administration. Samples were immediately frozen and stored at -80 °C or placed in 4% paraformaldehyde for fixation.

*RNA expression*

Harvested fetal lungs were frozen at -80°C until analysis. Total RNA was isolated using Trizol reagent following supplier recommended protocols. Purified RNA was quantified using a NanoDrop™ spectrophotometer and cDNA synthesis was performed with 500 ng quantified RNA (superscript VILO cDNA synthesis kit). qPCR experiments were conducted with SYBR™ Green Master Mix for 45 cycles (denaturation: 95 °C, annealing: 58 °C, extension: 72 °C) using the primer sequences reported in Table S6. Melt curve plots were used to determine target specificity of the primers. ΔΔCT method was used to determine normalized relative gene expression.

*Immunofluorescence*

Fetal rat lungs were fixed using 4% paraformaldehyde for 18 hours, paraffin-embedded, sectioned into 4 µm slices, and stained with primary antibodies reported in Table S7, as reported previously (*12*). A Leica SP8 lightning confocal microscope (Wetzlar, Germany) was used to image samples using the same laser power and exposure between conditions.

*Protein expression*

Protein from fetal rat lungs was isolated by incubation with cell extraction buffer supplemented with protease inhibitors, and sonication for 3 cycles of 10 seconds each. 20 µg of purified protein from each sample, quantified using the Pierce Bradford Assay, was probed for SPC and PDPN (Table S7). ß-Actin was used as a loading control.

*In vivo AFSC-EV tracking*

For tracking the location of AFSC-EVs following administration in vivo, a subset of experiments was conducted with either IA or IV injection of 100 µL of fluorescently labeled AFSC-EVs (ExoGlow™-Vivo) at E18.5. At E21.5, the whole body of the fetus or individually harvested organs were imaged using the IVIS® Spectrum In Vivo Imaging System – PerkinElmer (CFI Facility, University of Toronto). Briefly, whole fetuses were covered in talc powder to ensure optimal contrast and imaged with 2D fluorescence and 3D bioluminescence systems. Individual organs were reconstructed and imaged with 2D fluorescence using automatic exposure within an optimal range for quantification, avoiding overexposure during image acquisition, as recommended by the manufacturer. Saline-injected fetuses served as control. IVIS system was calibrated using 2 μL of ExoGlow™-Vivo. 3D reconstructions were generated using Living Image Software (version 4.7).

*Single nucleus RNA-sequencing experiments*

Nuclei were extracted from frozen lung tissue using Singulator TM 100 (S2 Genomics), with the following settings: nuclear reagent, S2; incubation temp, cold; mixing type, top; mixing speed, fastest; disruption type, default; disruption speed, medium. RNase inhibitor was added to the Singulator Cartridge to reach a final concentration of 0.2 U/μl. Nuclei collected from the Singulator was spun at 800 g, at 4 °C for 10 min. Supernatant was removed, following by resuspending nuclei in freshly made cold Wash and Resuspension Buffer (1X saline, 1% bovine serum albumin, 0.2 U/μl). The wash was repeated twice before proceeding into the 10X Genomics single-cell 3' v3.1 assay and processed as described by the protocol provided by 10X Genomics. Libraries construction and library sequencing were proceeded as described in the 10X Genomics protocol using NovaSeq 6000. FastQ files were obtained using Cell Ranger (10X Genomics, cellranger-6.0.0), aligned to the Rattus norvegicus version 6 genome, and quantified with Cell Ranger count function using default settings. Further downstream analysis was performed in R (version 4.0.2) and Seurat (version 4.0.3), with default settings unless specified otherwise. Data were analyzed either by replicate or by pooling reads by condition. Filtering of data was performed to remove cells with raw data was log normalized and scaled using default values. Variable features and principal components were then calculated using default values. Ribosomal genes were removed with pattern = "^Rp[sl][[:digit:]]|^Rplp[[:digit:]]|^Rpsa" from normalized data. UMAP dimensionality reductions were performed with default values. Batch effects were not evident in the dimensionality reductions, and therefore, the data was then analyzed as-is without further corrections. For each major cell type, we performed differential expression analysis using Seurat with MAST method between different conditions (e.g., Control+saline vs CDH+saline). We used Benjamini Hochberg corrected p-values for assessing statistical significance. Genes with p-value < 0.05 were considered as differentially expressed. Ligand-receptor analysis was performed using CellChat (Version 1.1.3) with default parameters.

*miRNA-mRNA regulatory network analysis*

Our previously published dataset of AFSC-EV miRNA cargo (*12*) was used to interrogate possible regulatory networks that were downregulated following AFSC-EV administration. We used miRNAs that were overexpressed in AFSC-EV cargo compared to another source of EVs from mesenchymal stromal/stem cells (*12*). Significantly downregulated genes in CDH+AFSC-EVs vs. CDH+saline groups were considered targets of AFSC-EV cargo miRNA. MultiMiR (v1.16.0) was used to link miRNAs from predicted and validated databases (miRecords, miRTarBase, TarBase) with targets. miRNA-mRNA pairs were visualized and shown as an interaction network generated using Cytoscape (3.8.2).

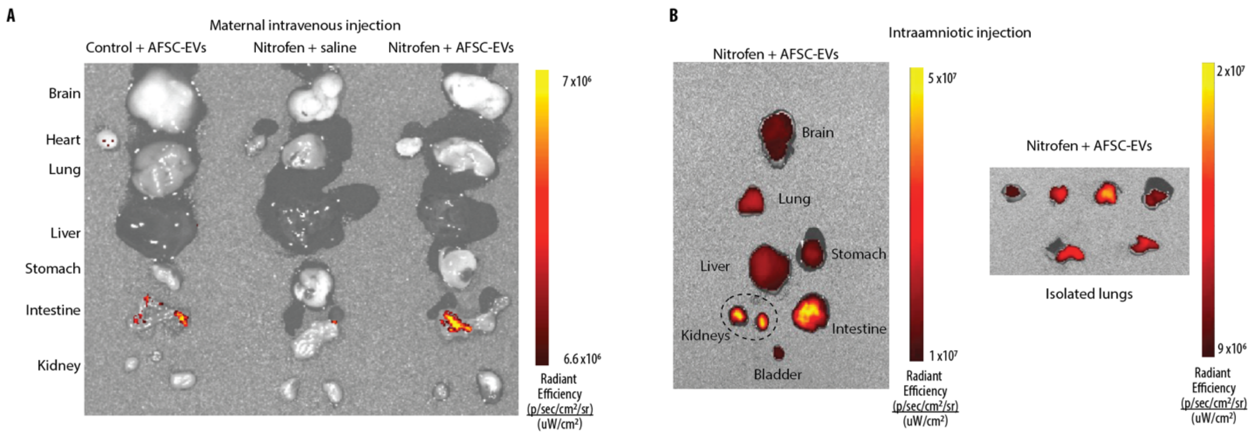


**fig. S1. In vivo effects on individual fetal organs.** (**A**) Three representative 2-dimensional bioluminescence IVIS images of organs harvested from embryonic day E21.5 fetal rats following maternal intravenous injection of ExoGlowVivo-stained AFSC-EVs in control pups (left), or pups exposed to nitrofen and treated with AFSC-EVs (right), compared to nitrofen-exposed pups that received saline alone (middle). Control+saline (n=3), Nitrofen+saline (n=3), Nitrofen+AFSC-EVs (n=3). (**B**) Representative bioluminescence IVIS image of a nitrofen-exposed fetus treated with ExoGlowVivo-stained AFSC-EVs via intra-amniotic injection. Left side indicates all organs (n=10 biological replicates), right side shows fetal lungs from six different biological replicates.

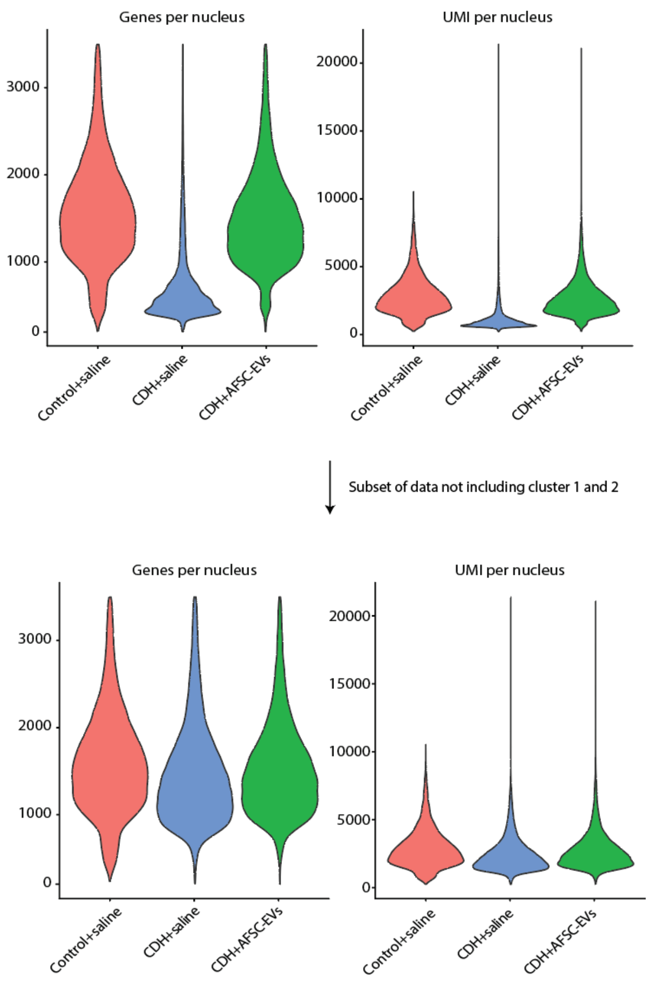


**fig. S2. Quality control metrics of single nucleus RNA-sequencing data.** Plots of genes per nucleus and unique molecular identifiers (UMI) per nucleus in the final dataset (top) and subset of data that did not include cluster 1 and 2 from **Fig. 5F** (bottom).

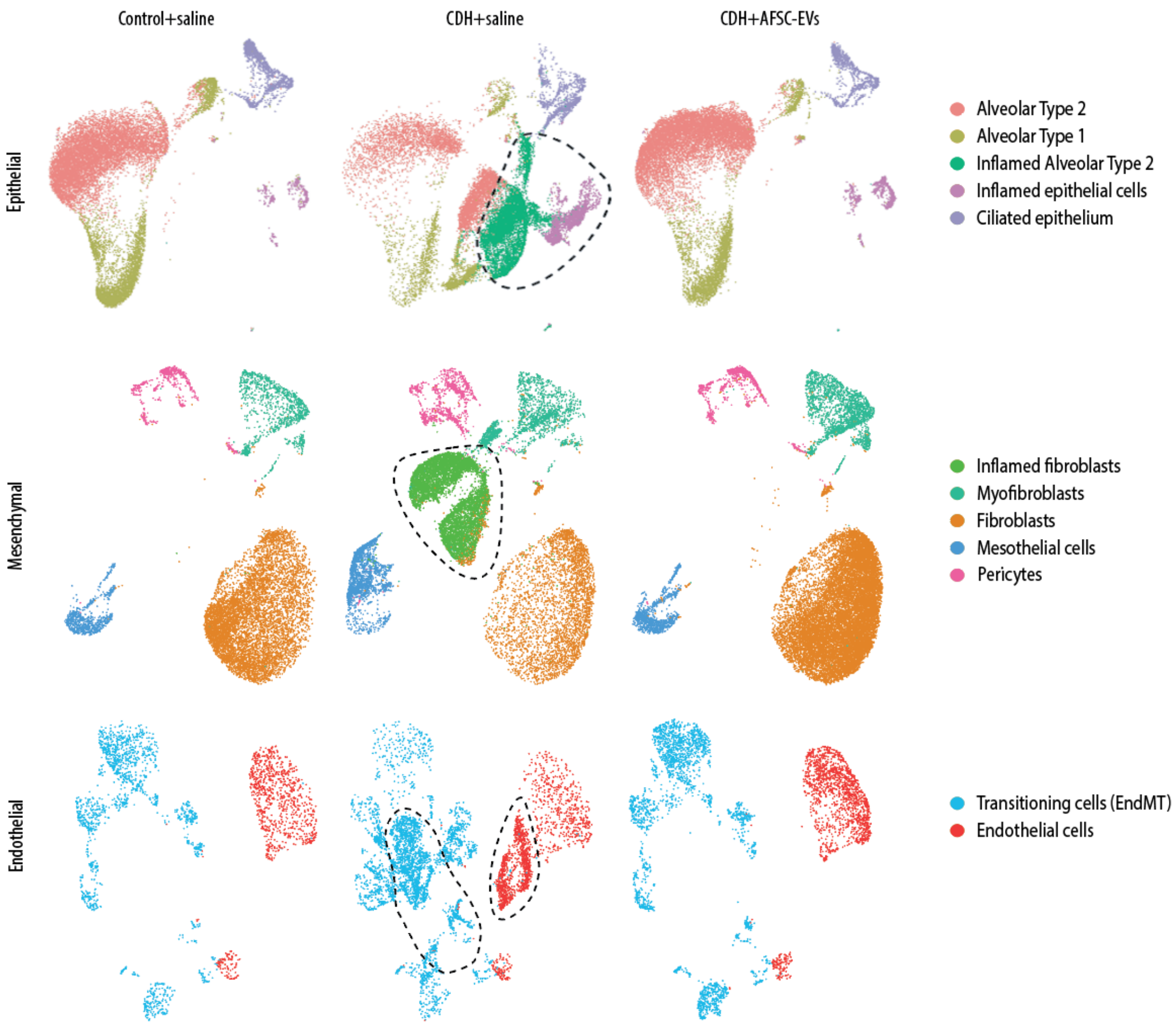


**fig. S3. UMAP depicting major cell types split by condition.** Individual UMAPs of epithelial (top), mesenchymal (middle), and endothelial (bottom) cell types split by conditions: Control+saline (left), CDH+saline (middle), and CDH+AFSC-EVs (right). Colors indicate subgroups. Dotted lines outline the nuclei that are uniquely expressed in CDH+saline lungs.

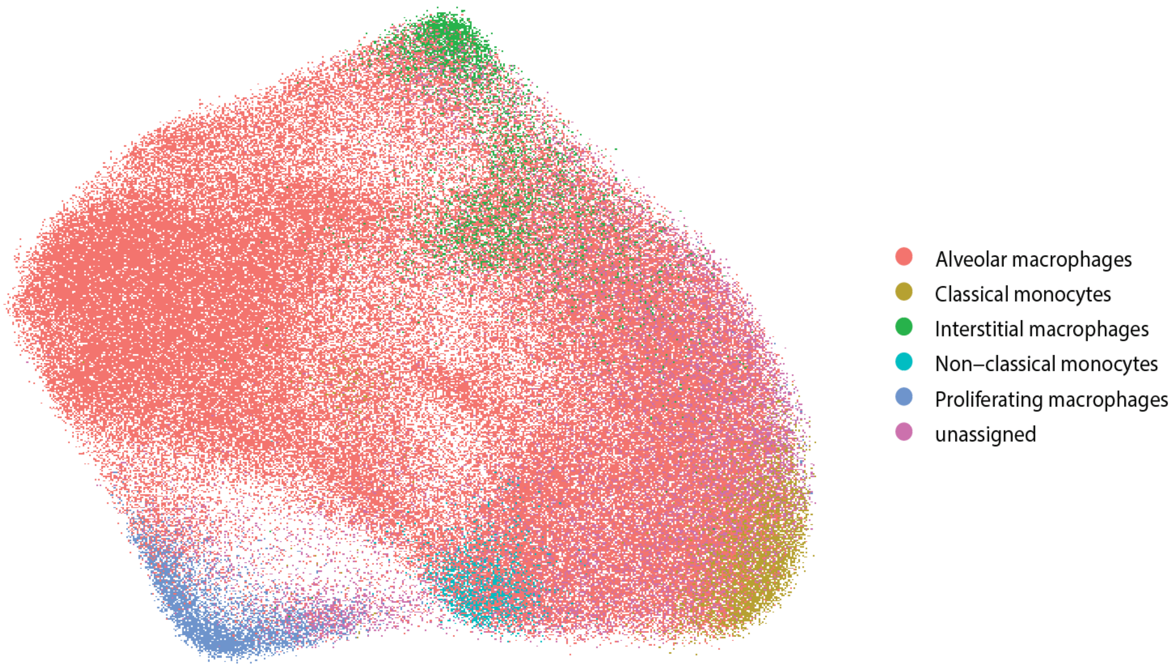


**fig. S4. Global UMAP of predicted subtypes in cluster 1 and 2.** ScPred tool was used to predict subtypes using machine learning on adult rat lung single cell data (*54*, *55*). Colors indicate subgroups.

**Table S1: Number of nuclei in each cluster for snRNA-sequencing data analysis**

| | | number of nuclei per cluster | | |
|---|---|---|---|---|
| **Name** | **cluster number** | **Control + saline** | **CDH + saline** | **CDH + AFSC-EVs** |
| Macrophage group 1 | 1 | 0 | 91109 | 0 |
| Macrophage group 2 | 2 | 247 | 89187 | 739 |
| Alveolar type 2 | 3 | 9721 | 5248 | 10578 |
| Fibroblast | 4 | 7045 | 3705 | 14069 |
| Immune | 5 | 1783 | 6610 | 4165 |
| Alveolar type 1 | 6 | 4526 | 2459 | 3352 |
| Inflamed fibroblast | 7 | 6 | 6931 | 3 |
| Inflamed alveolar type 2 | 8 | 13 | 6843 | 21 |
| Myofibroblast | 9 | 1150 | 1794 | 2185 |
| EndMT | 10 | 1575 | 3629 | 2386 |
| Endothelial | 11 | 974 | 1731 | 1614 |
| Mesothelial | 12 | 858 | 2114 | 1138 |
| Ciliated epithelial | 13 | 1417 | 1109 | 891 |
| Inflamed epithelial | 14 | 366 | 1864 | 880 |
| Pericytes | 15 | 630 | 1077 | 911 |

**Table S2: Genes upregulated in fetal lungs following in vivo AFSC-EV administration**

| Gene | Name | Function | Role during lung development and CDH (if known) | References |
|---|---|---|---|---|
| ***Calcrl*** | Calcitonin receptor-like | Receptor of adrenomedulin (AM), which plays a key role in endothelial growth and development. AM is ubiquitously expressed, including in blood vessels and lungs. | AM has potential anti-inflammatory, antioxidant, angiogenic, and vasodilatory properties in the lungs. AM was found to increase pulmonary angiogenesis and attenuate alveolar simplification and pulmonary hypertension in a rat model of hyperoxia-induced BPD. | (*122-124*) |
| ***c-Kit*** | Tyrosine-protein kinase KIT, CD117 | Proto-oncogene that interacts with cell growth factors and plays a role in cell survival, multiplication, and differentiation. | In the lung, c-Kit is responsible for maintenance of normal alveolar architecture, regulation of epithelial cell clonal expansion, and vascular formation. | (*125*, *126*) |
| ***Igf*** | Insulin-like growth factor | Protein-coding gene that is involved in mediating growth and development in many cells and tissues. | The Igf system plays a pivotal role in the development and differentiation of the fetal lung. IGF receptor type 1 and type 2 are downregulated in nitrofen-induced hypoplastic lungs. | (*127-129*) |
| ***Vegf*** | Vascular endothelial growth factor | Potent mitogenic and angiogenic factor that is responsible for differentiation and proliferation of endothelial cells during embryogenesis. | Vegf plays a key role in the development of the pulmonary capillary bed. Experimental studies have suggested that changes in the Vegf signaling pathway are associated with pulmonary hypoplasia in CDH. | (*130-132*) |
| ***Fgf*** | Fibroblast growth factor | Regulates a broad spectrum of biological functions, including cellular proliferation, survival, migration and differentiation. | Fgf signaling plays a key role during lung development, injury, and repair. | (*133*, *134*) |
| ***Egf*** | Epidermal growth factor | Plays an important role in the growth, proliferation and differentiation of numerous cell types. | Egf enhances alveolar type II cell differentiation and stimulates surfactant protein A synthesis. Its receptor is found in the alveolar epithelium during differentiation, suggesting an important role for Egf during human fetal lung development. Exogenous EGF | (*135*, *136*) |

| | | | | |
|---|---|---|---|---|
| | | | improves lung growth in the nitrofen-induced CDH model. | |
| ***Gas*** | Growth arrest-specific protein 1 | Protein-coding gene that plays a key role in processes related to proliferation, inflammation, tissue repair, and angiogenesis. | In the lung, Gas signaling has been shown to reduce alveolar inflammation and acute ischemia-reperfusion injury. Furthermore, Gas is a promising molecular marker for therapies of chronic thromboembolic pulmonary hypertension. | (*137*, *138*) |
| ***Pros*** | Protein-S1 encoding gene | Vitamin-K-dependent plasma protein-coding gene with anticoagulant properties. | Pros1 signaling regulates lung cancer cell proliferation, migration, and angiogenesis. | (*139*, *140*) |
| ***Sema3*** | Semaphorin 3A gene | Protein-coding gene with multifunctional roles in embryonic development, immune regulation, and vascularization. | Sema3a stimulates branching morphogenesis and cell proliferation. | (*141*, *142*) |
| ***Nrg*** | Neuroregulin 1 | Belongs to the EGF family and is involved in tissue development and maturation. | Nrg controls fetal lung maturation through mesenchymal-epithelial interactions. Nrg1 induces branching morphogenesis in the developing lung through a P13K signal pathway. CDH fetal lungs are associated with decreased NGR1 expression in the lamb model of CDH. | (*143-145*) |
| ***Csf*** | Colony stimulating factor 1 | Protein-coding gene that controls the production, differentiation, and function of macrophages. | CSF1 has a pivotal role in fetal lung development, with regenerative effects through polarization of macrophages towards an M2 phenotype. | (*146*) |

**Table S3: Predicted cell types from cluster 1 and 2 of CDH+saline lungs**

| | number of nuclei | percentage |
|---|---|---|
| Alveolar macrophages | 140382 | 77% |
| Classical monocytes | 5879 | 3% |
| Interstitial macrophages | 6511 | 4% |
| Non-classical monocytes | 2287 | 1% |
| Proliferating macrophages | 4942 | 3% |
| Unassigned | 21281 | 12% |

**Table S4: Number of nuclei within each condition of predicted cell types from cluster 5 (immune cells).**

| Cell Subtypes | Control + saline | CDH + saline | CDH + AFSC-EVs |
|---|---|---|---|
| Alveolar macrophages | 53 | 168 | 79 |
| B cells | 113 | 147 | 268 |
| CD8 T cells | 12 | 4 | 18 |
| Classical monocytes | 338 | 939 | 983 |
| Conventional dendritic | 35 | 48 | 99 |
| ILC2 | 4 | 2 | 14 |
| Interstitial macrophages | 245 | 497 | 740 |
| Mast cells | 41 | 8 | 40 |
| Naive T cells | 384 | 161 | 518 |
| Neutrophils | 58 | 3506 | 182 |
| NK cells 1 | 37 | 24 | 65 |
| NK cells 2 | 16 | 10 | 41 |
| Non-classical monocytes | 25 | 161 | 72 |
| Plasmacytoid dendritic | 19 | 9 | 33 |
| Proliferating macrophages | 55 | 314 | 243 |
| Proliferating T cells | 33 | 42 | 98 |
| Regulatory T cells | 18 | 19 | 57 |
| unassigned | 297 | 551 | 615 |

**Table S5: Details from human lung autopsy specimens used in this study.**

| Case # | CDH | Age (Gestational weeks) | Description |
|---|---|---|---|
| 1 | + | 19 | Left-sided CDH, pulmonary hypoplasia |
| 2 | + | 20 | Left-sided CDH, pulmonary hypoplasia |
| 3 | + | 26 | Absence of left diaphragm, pulmonary hypoplasia |
| 4 | + | 27 | Left-sided CDH, pulmonary hypoplasia, intrauterine growth restriction, stillborn |
| 5 | - | 19 | Elected termination of pregnancy |
| 6 | - | 20 | Elected termination of pregnancy |
| 7 | - | 18.4 | Elected termination of pregnancy |
| 8 | - | 19 | Elected termination of pregnancy |

**Table S6. Primer sequences used in this study for rat qPCR.**

| Target | Forward primer | Reverse primer |
|---|---|---|
| *Fgf10* | 5’-AGCTGTTCTCCTTCACCAAGTA-3’ | 5’-ACTCCGATTTCCACTGATGTTA-3’ |
| *Pdpn* | 5’-CCTCCACTTGCCAGCAGTA-3' | 5’-GCATGTGGTCCTCAATCATAAC-3’ |
| *Sftpc* | 5’-AACGCCTTCTCATCGTGGTT-3' | 5’-GGCTTATAGGCGGTCAGGAG-3’ |
| *Sftpa* | 5’-AACGAGGCCATTGCAAGTATT-3' | 5’-GAAGCCCCATCCAGGTAGT-3' |
| *Gapdh* | 5' -AGTGCCAGCCTCGTCTCATA- 3' | 5' -GAGAAGGCAGCCCTGGTAAC- 3' |

**Table S7: Details of antibodies used in this study.**

| Target | Antibody | Company | Immunofluorescence concentration | Western blotting concentration |
|---|---|---|---|---|
| SPC | ab40879 | Abcam | 1:200 | - |
| SPC | AB3786 | EDM Millipore | - | 1:500 |
| PDPN | P5374 | ThermoFisher | 1:200 | 1:500 |
| ß-actin | ab8226 | Abcam | - | 1:1000 |
| CD68 | ab283654 | Abcam | 1:100 | - |
| pNFκB-p65 (Ser 536) | #3036 | Cell signal | 1:100 | - |
| TNFα | sc-1350 | Santa Cruz | 1:200 | - |
