## Supplementary figures and images for "Administration of amniotic fluid stem cell extracellular vesicles promotes development of fetal hypoplastic lungs by immunomodulating lung macrophages"

### Data file S1

#### Slide 1
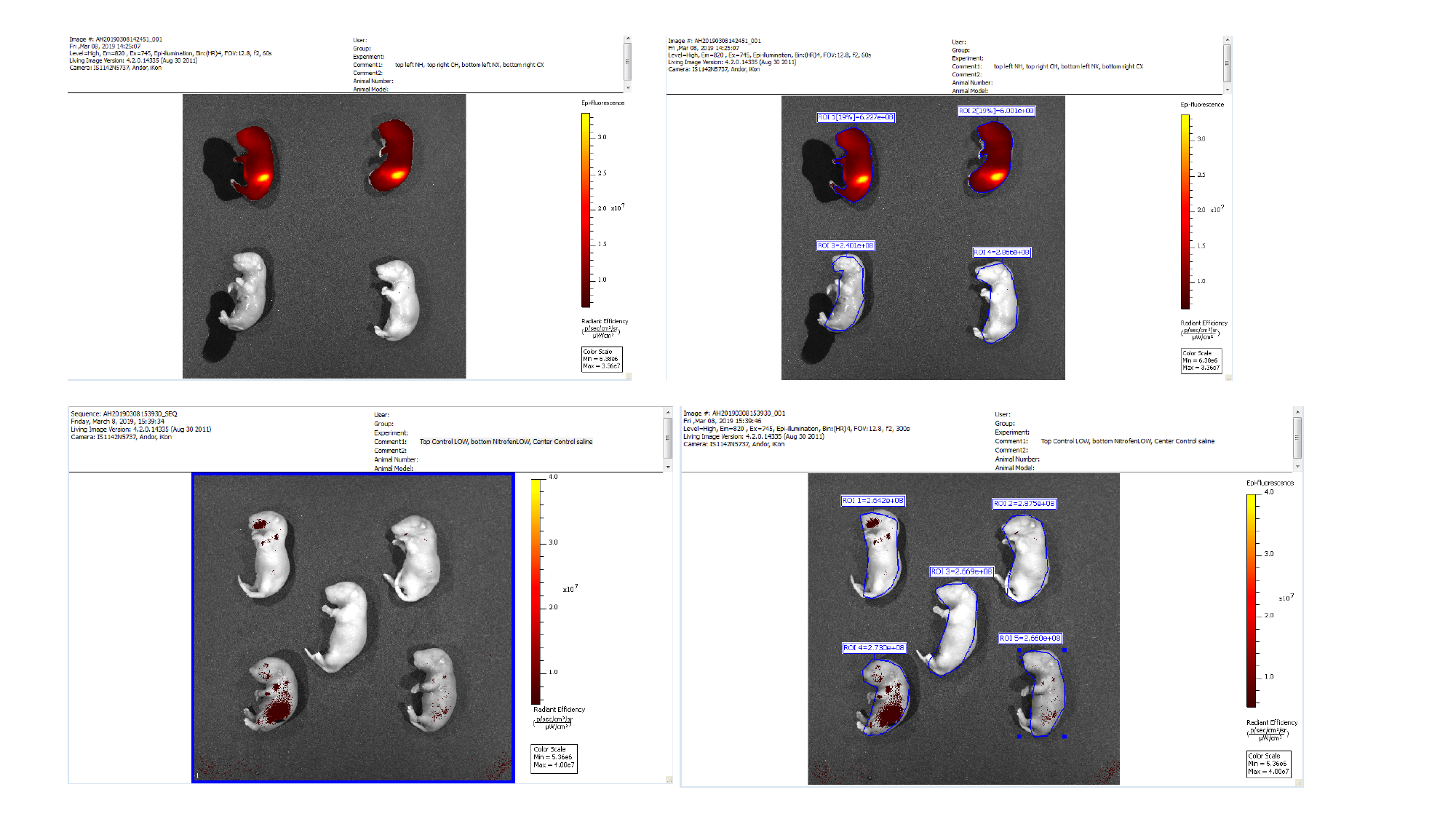

#### Slide 2
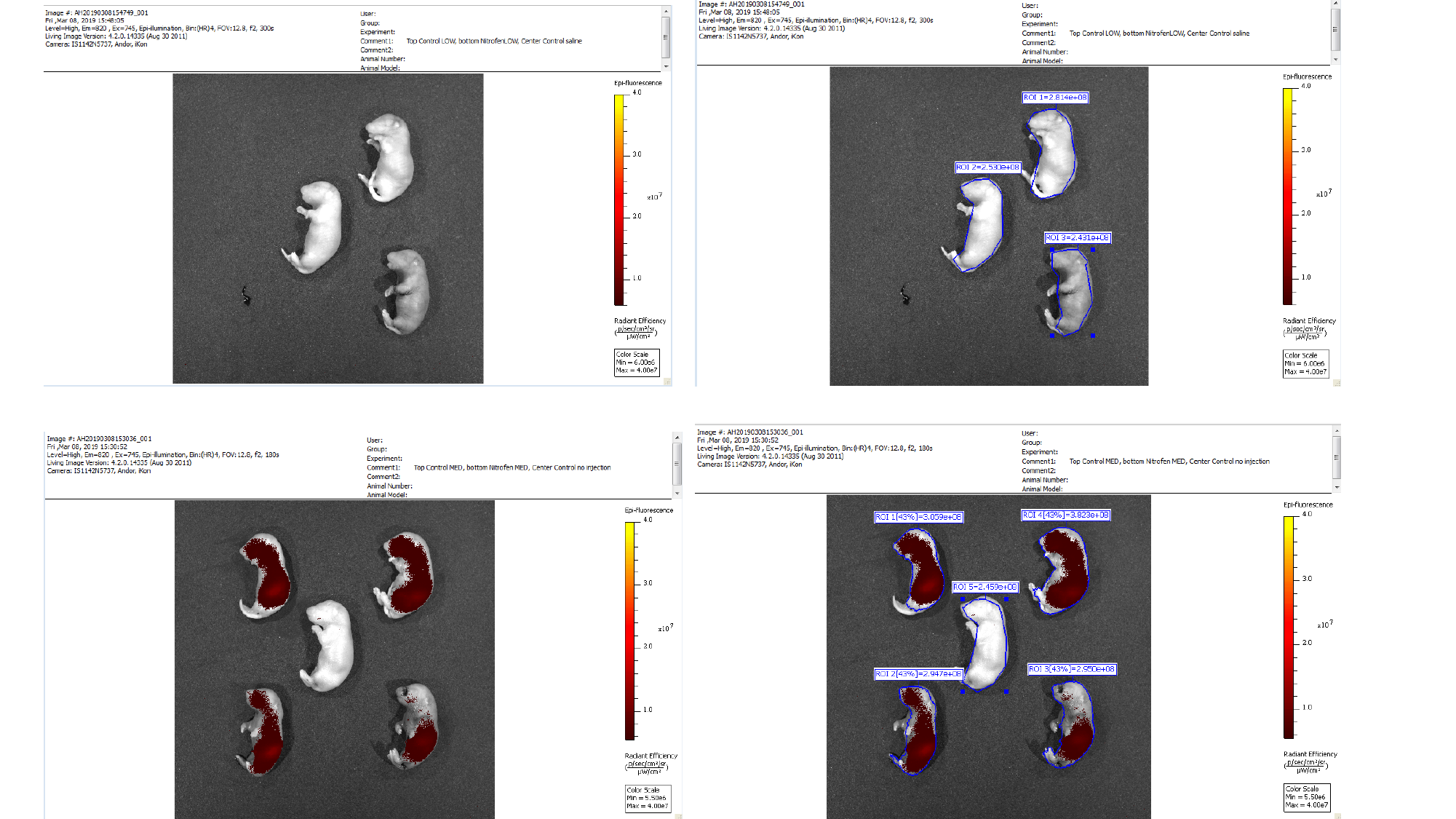

#### Slide 3
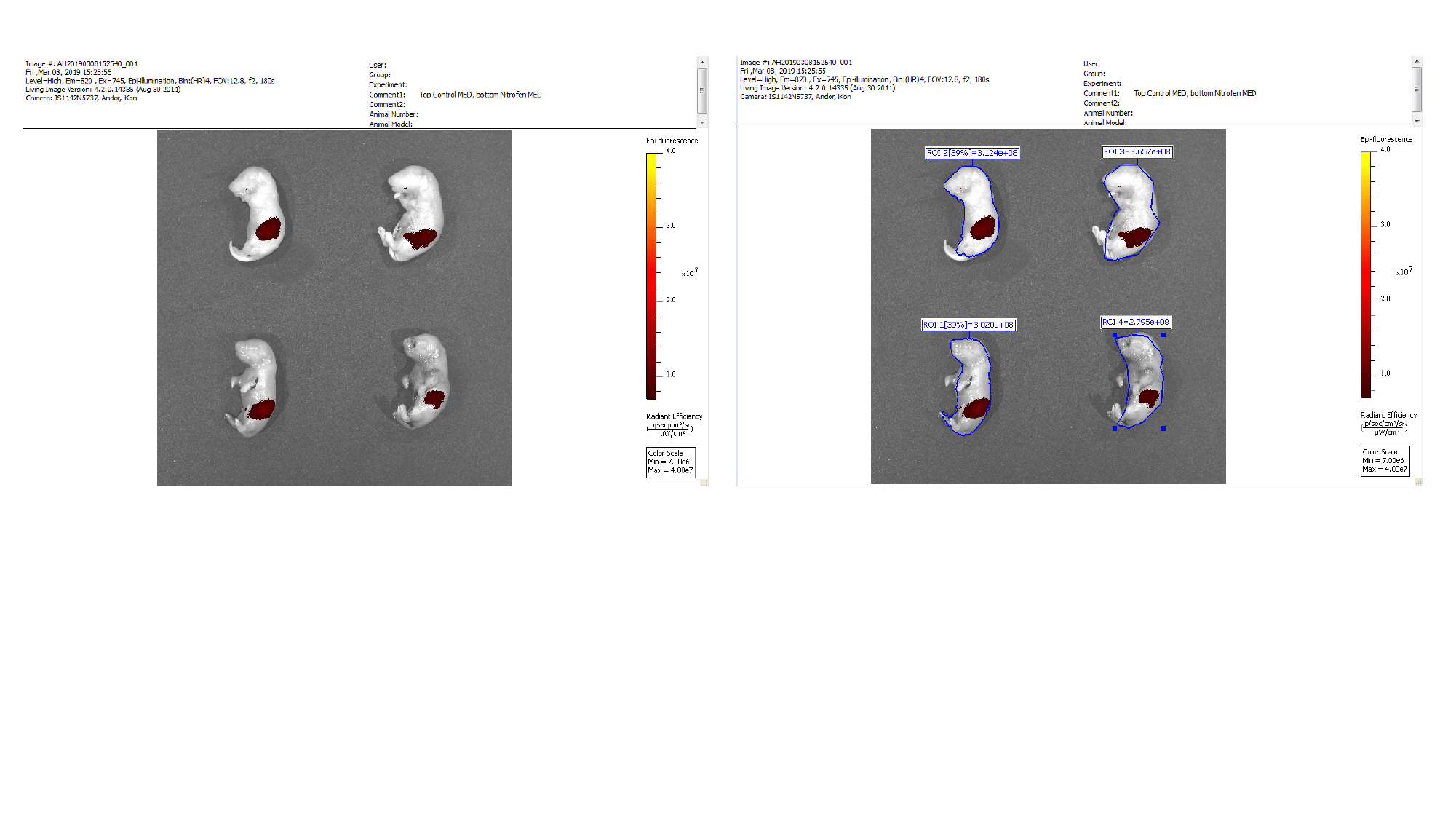

### Data file S2

#### Slide 1
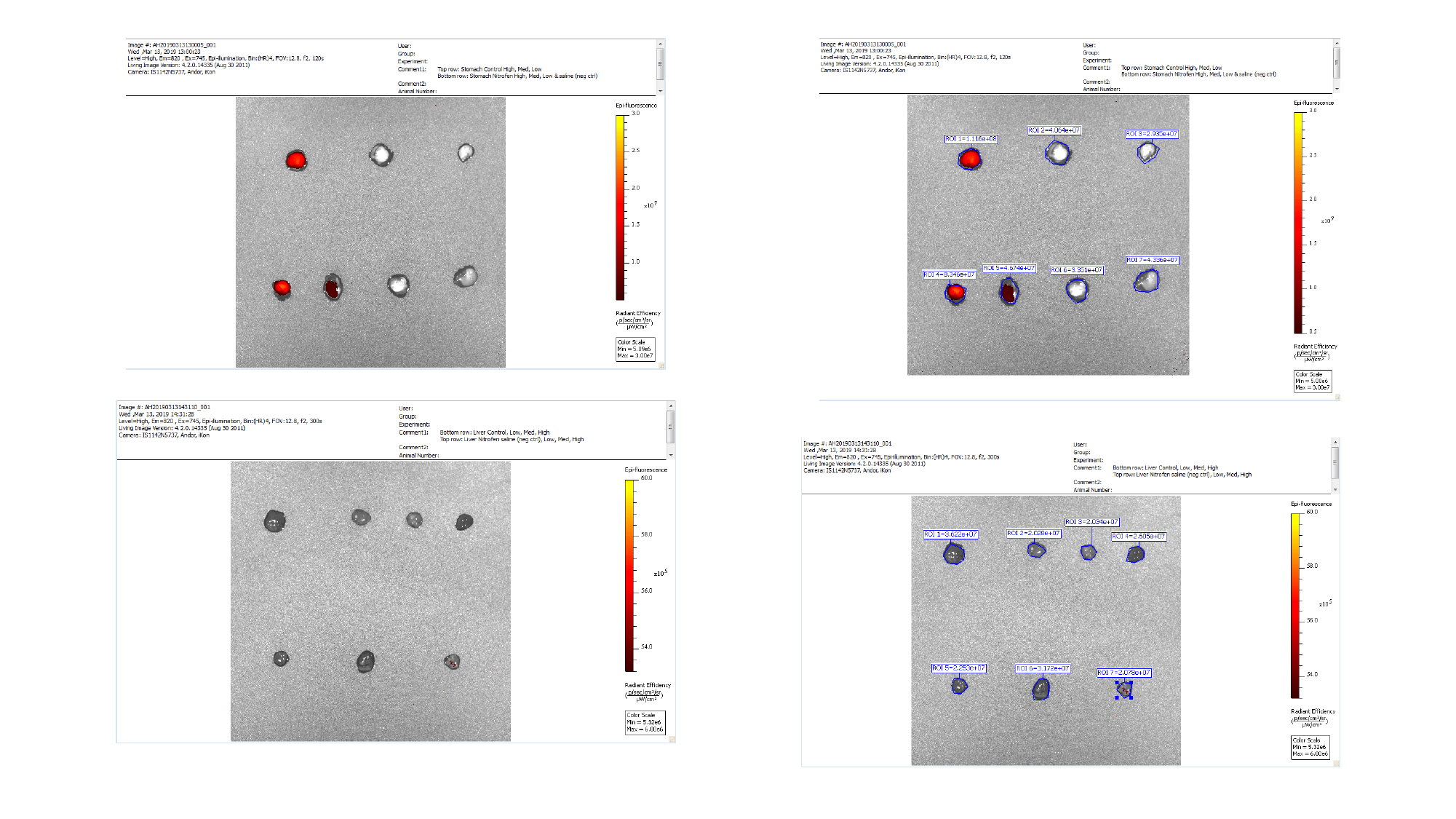

#### Slide 2
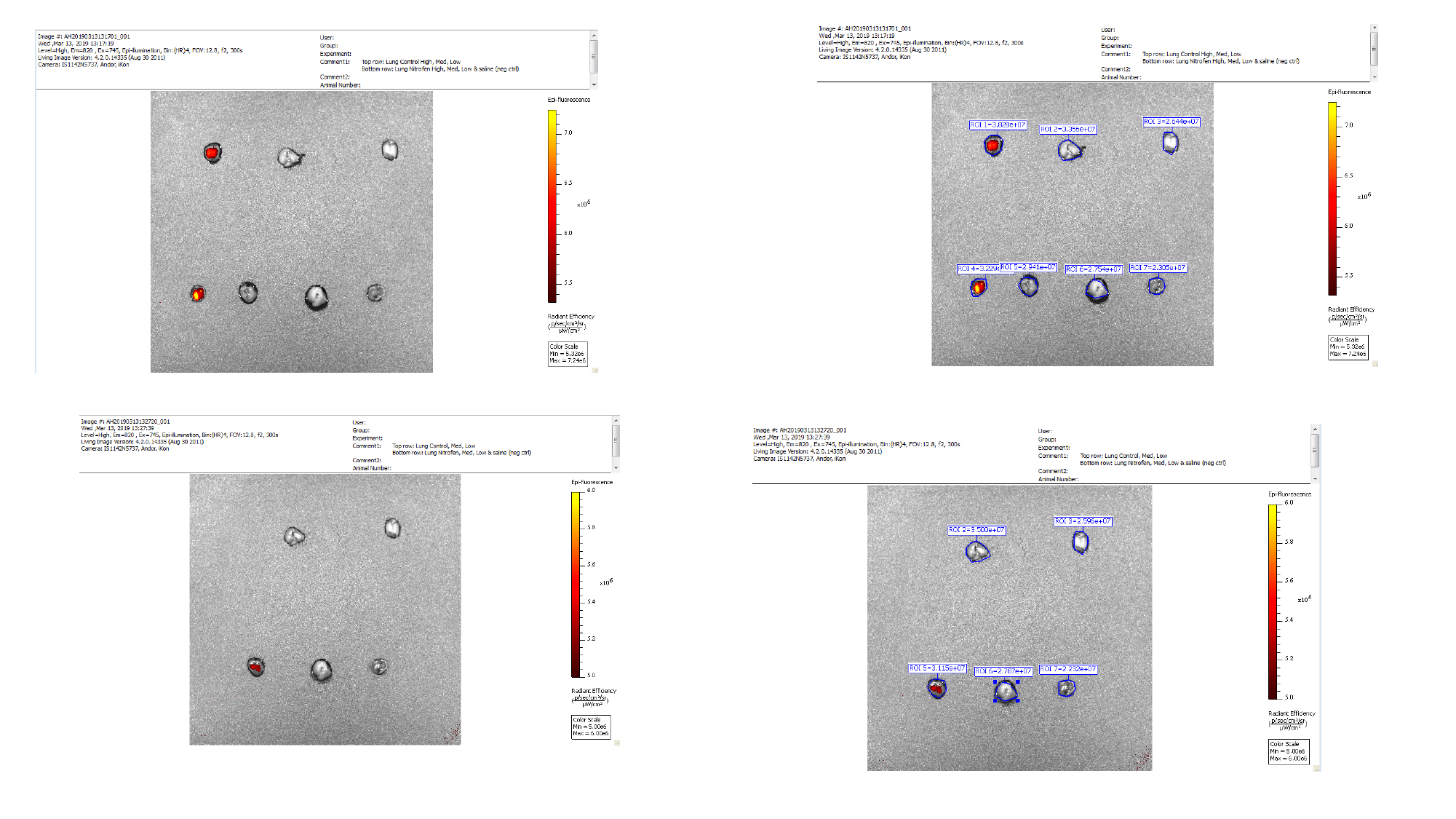

#### Slide 3
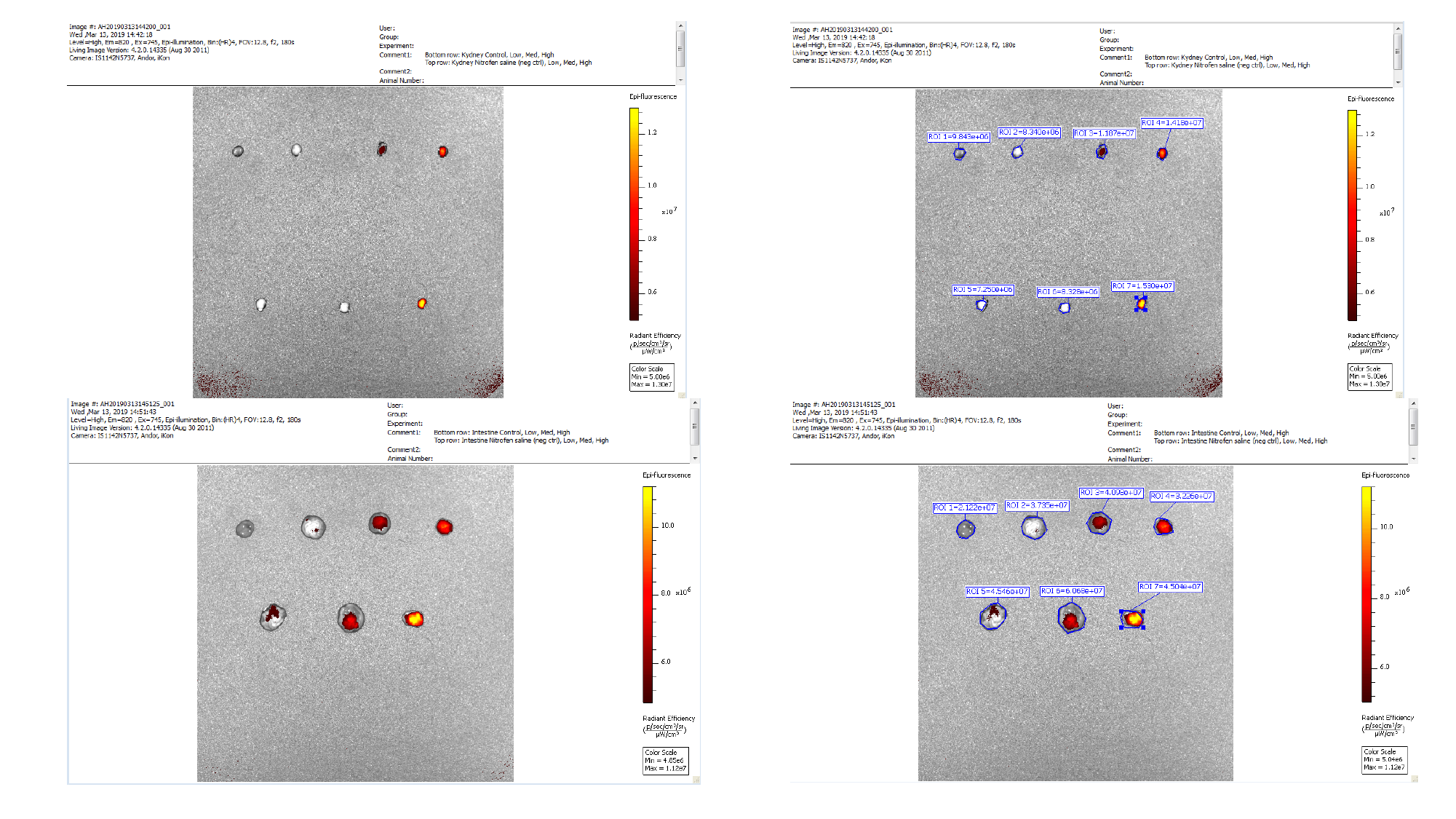

#### Slide 4
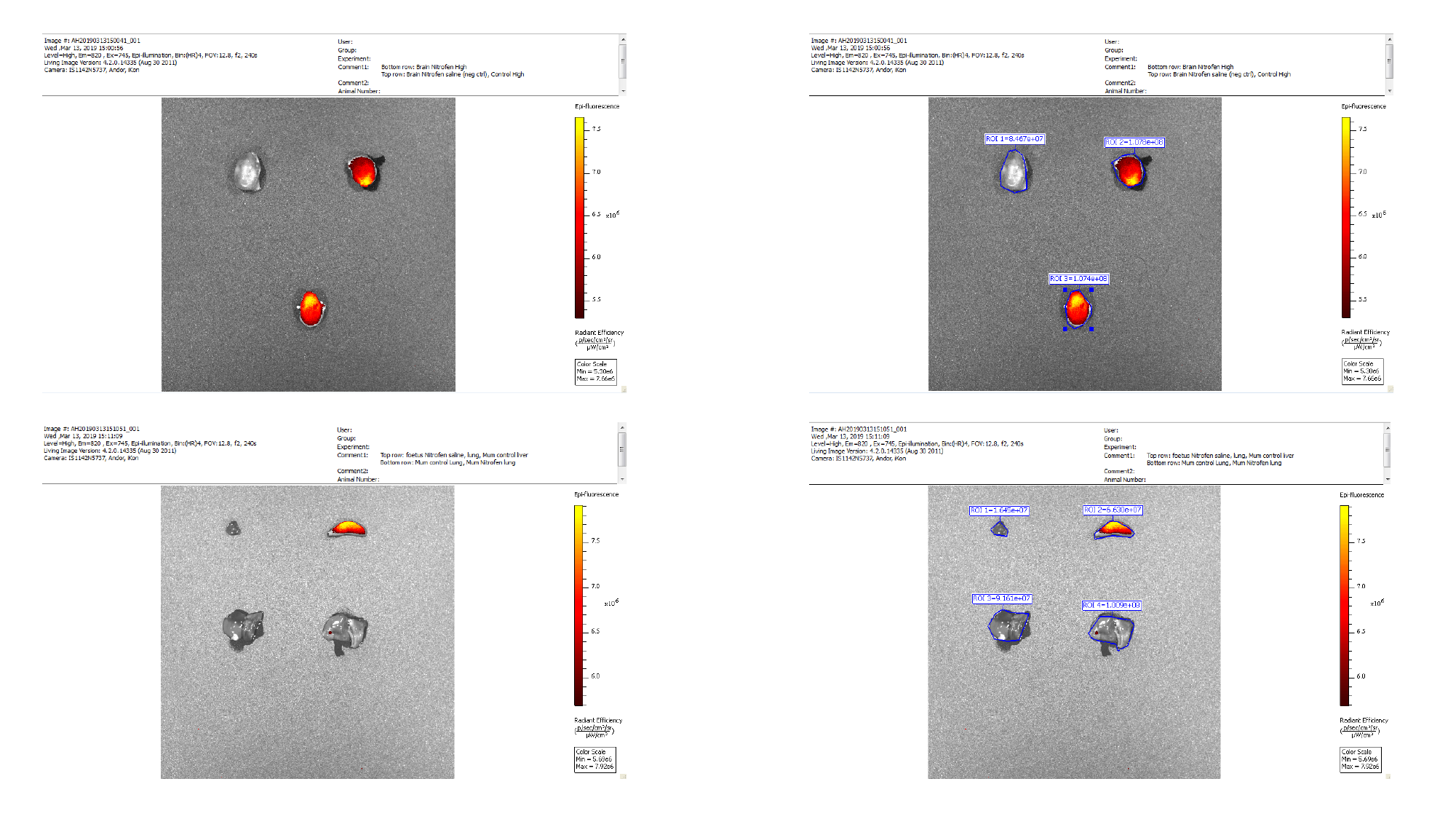

#### Slide 5
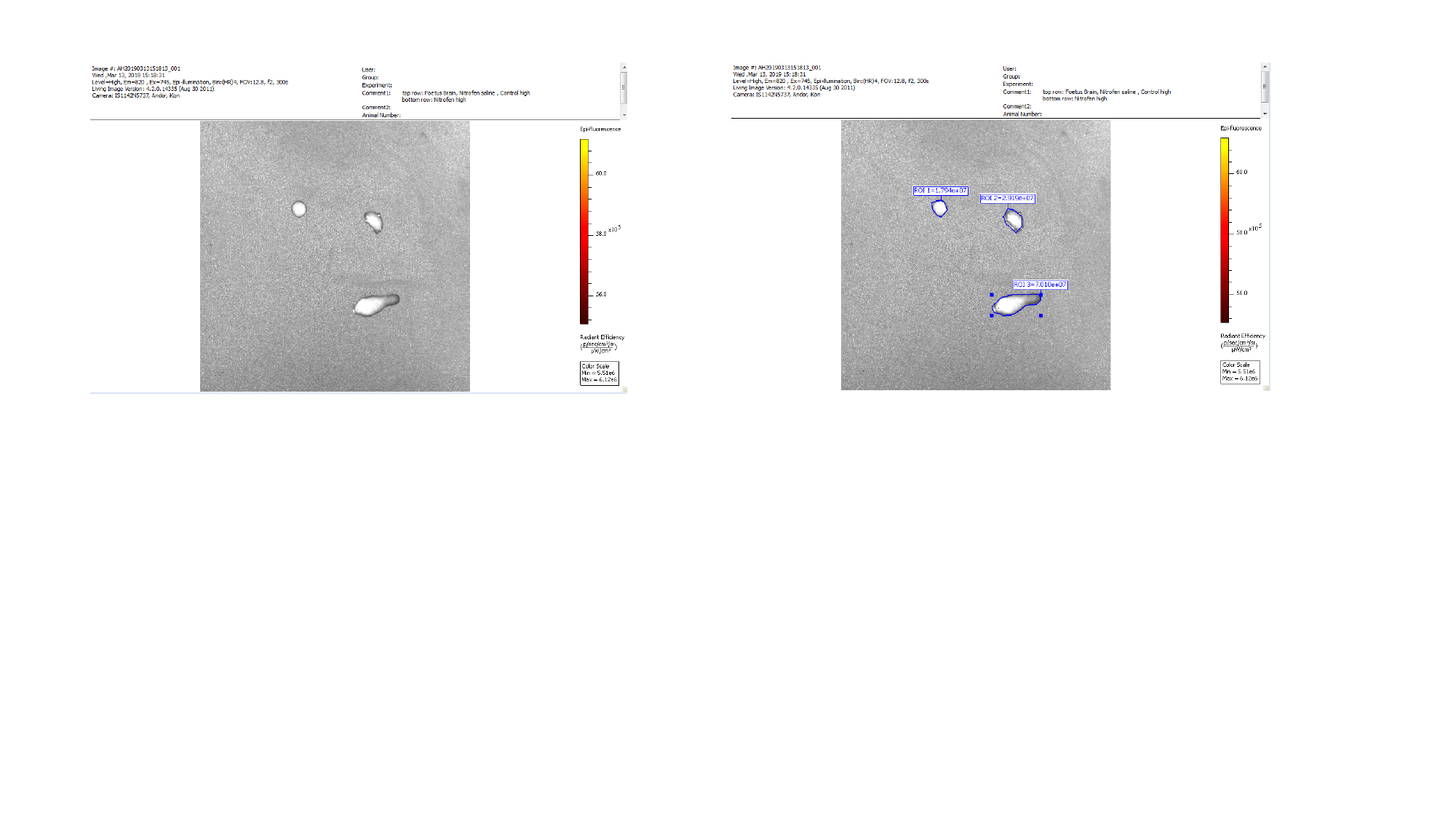
